# Super-pangenome analysis of 3562 human and animal papillomavirus isolates illuminates their genome and pathogenicity evolution

**DOI:** 10.1101/2025.07.20.664904

**Authors:** Huihui Li, Jingtao Chen, Xinxin Zhang, Xing Zhang, Ludong Yang, Yi Ding, Congying Chen, Yan Shuai, Ming Song, Jihong Liu, Lin Feng, Jing Li, Jia-Xing Yue

**Author notes:** corresponding authors. (JL); (JXY). contribute equally (Huihui Li, Jingtao Chen, Xinxin Zhang).

## Abstract

Papillomaviruses (PVs) are groups of ubiquitous DNA viruses infecting a wide range of vertebrate hosts, including human. The infection of some human PVs (i.e., HPVs) are strongly associated with cancers, but it remains elusive how such pathogenicity evolved at the genomic level. Here we performed a super-pangenome analysis on a compendium of well-curated 3562 human and animal PV genomes. We found their global genomic diversity being strongly associated with their body sites and tissue origins, but not with their host, geographical, or disease specificity. By reconciling time-resolved phylogenies of PVs and their hosts, we revealed a high-resolution view on how virus-host cospeciation, horizontal host switches, and duplication/loss turnovers collectively shaped the evolution of PVs. We identified key selection-driven mutations that defined the genus-specific divergence of PVs, with those underlying the rise of Alpha-PVs highly enriched in the N-terminal domain of *E1* gene. Especially, we identified a group of risk-defining variant sites showing contrasting genotypes between low-risk and high-risk HPVs. With four independent PV genome cohorts, we further validated their high sensitivity and specificity on HPV risk classification even at single site level, underscoring their application values in large-scale HPV screening tests. Finally, with cervical and oropharyngeal cancer patient cohorts assembled by this study, we also evaluated the prognostic value of specific HPV16 variants in predicting patient survival. Taken together, such illumination on the genome and pathogenicity evolution of PVs paves the road for better prevention, control, and treatment of PV-related cancers.

**Significance statement:** Papillomaviruses (PVs) are ancient and widespread DNA viruses co-evolved with many vertebrate hosts, including human. Some human PVs (e.g., HPV16 and HPV18) bear strong association with cancers, raising a pressing public health challenge. A better understanding of how such pathogenicity evolved holds the key in addressing this challenge. Here we systematically investigated the genomic and evolutionary diversity of PVs via a pangenome study on over 3500 human and animal PV isolates, which revealed key selection-driven mutations shaping their genome and pathogenicity evolution. With independently acquired HPV-associated cancer cohorts, we further demonstrated how some of these important HPV variants can serve as clinically informative predictive and prognostic biomarkers. Such comprehensive characterization on PVs’ genome diversity, pathogenicity evolution, and clinical implication paves the road for better prevention, control, and treatment of PV-related cancers.

## Introduction

Papillomaviruses (PVs) is a ubiquitous group of double-stranded DNA viruses that infect the basal keratinocytes of cutaneous and mucosal epithelia in a wide range of vertebrate host species such as fish, reptiles, birds, and mammals, including human[1]. A typical PV genome has a size of ∼5–8 kb and encodes 6–9 genes[2], with genes *E5* and *E6* considered as oncogenes[3–5]. Traditionally, PVs are classified into different taxonomic groups (e.g., genera, species, and types) according to the nucleotide sequence divergence of their major capsid encoding-gene *L1*[6,7]. Members from each genus share >60% *L1* nucleotide sequence identity, while different species within a genus share 60%–70% sequence identity. A novel PV type is assigned if it shows <90% similarity to any other previously defined PV types. So far, >220 types have been defined for human PVs (i.e., HPVs), which distributed across five genera: Alphapapillomavirus (Alpha-PV), Betapapillomavirus (Beta-PV), Gammapapillomavirus (Gamma-PV), Mupapillomavirus (Mu-PV), and Nupapillomavirus (Nu-PV). Interestingly, HPV types that infect mucosal epithelium was only found in Alpha-PV, whereas HPVs types that infect cutaneous epithelium ubiquitously distributed across all of these five genera[1]. In comparison, PVs from animal hosts distribute across more different genera (e.g., Omicron, Upisilon, Omega, etc.) as reflected by their larger genomic divergence.

In terms of pathogenicity, many PV types are either completely asymptomatic or considered as low-risk (e.g., HPV types 6, 11, 40, 42, 43, 44, 54, 61, 70, 72, 81, and CP6108) as they only cause benign tumors such as warts or papilloma. In contrast, some other PVs types are classified as high-risk (e.g., HPV types 16, 18, 31, 33, 35, 39, 45, 51, 52, 56, 58, 59, 66, 68, 73, and 82) or probably high-risk (e.g., HPV types 26, 53, and 66) due to their association with human cancers[6–8]. For example, it has been estimated that ∼90% of cervical cancers and ∼70% of oropharyngeal cancers are associated with HPV infections[9]. Therefore, a better understanding of the carcinogenicity of HPVs holds the key in the control and treatment of HPV-associated cancers.

Given the critical clinical importance of HPVs, genome sequencing and comparison have become a powerful approach to decode their genomic diversity in both human and non-human hosts. With this approach, previous studies suggested a general co-divergence relationship between PVs and their hosts in evolution (known as “Fahrenholz’s rule”) but also noticed some exceptions[10–13]. For example, PVs from five well-diverged genera (Alpha- to Nu-PVs) were all found in human, while some non-human primate PVs and HPVs are phylogenetically intermingled[14,15]. To reconcile such observation, hypothesis of niche-specific adaptation within the host was proposed, suggesting PV as an ancestral generalist gradually diverged and become niche-specific specialists in their recent evolution with primate hosts[12,14,16]. In addition, sporadic cases of horizontal host transfer has also been reported, probably occurred via cross-species infection[17].

To date, there have been thousands of PV genomes available in online databases such as Genbank, with hundreds of representative ones further curated in dedicated database such as PaVE[18]. Therefore, a super-pangenome-based study that covers all currently known PV lineages isolated from both human and animal hosts is expected to provide a comprehensive view on the global genomic and ecological diversity of PVs and shed light on their host-associated evolutionary history. In this study, we systematically curated and annotated a compendium of 3562 PV genomes with diverse host species origins (e.g., human, non-human primates, non-primate mammals, reptiles, birds, and fish) to derive a high-quality super-pangenome collection for PVs. With this super-pangenome, we characterize the genomic diversity of PVs in relation to their host specificity, geographic distribution, atomical tissue origin, risk and disease association, gene repertoire, and phylogenetic clustering. More importantly, we identified key genomic variants in PV genomes underlying their genus-specific diversification and pathogenicity evolution. By further incorporating multiple independent cohorts, we demonstrated the medical relevance and application value of some key HPV variants for the precision prevention and treatment of HPV-associated cancers.

## Results

### A super-pangenome view on the global diversity of PVs

To capture and understand the global genomic and ecological diversity of PVs, we collected, curated, and annotated a super-pangenome collection of 3562 PV genomes via both database inquiry and literature mining (Supplementary Table 1). Thorough metadata curation was also performed regarding their host species, geographic distribution, anatomic body site and tissue origins, risk and disease associations. The 3562 PVs include 3329 HPVs, 30 non-human primate PVs, 172 non-primate mammal PVs, 6 reptile PVs, 11 bird PVs, and 14 fish PVs, covering all known host species of PVs (Figure 1A). The 3329 HPVs collected here were isolated from the global population (Figure 1B), with most of them classified into the Alpha (84.9%), Beta (2.8%) and Gamma (12.2%) genera.

**Figure 1.**
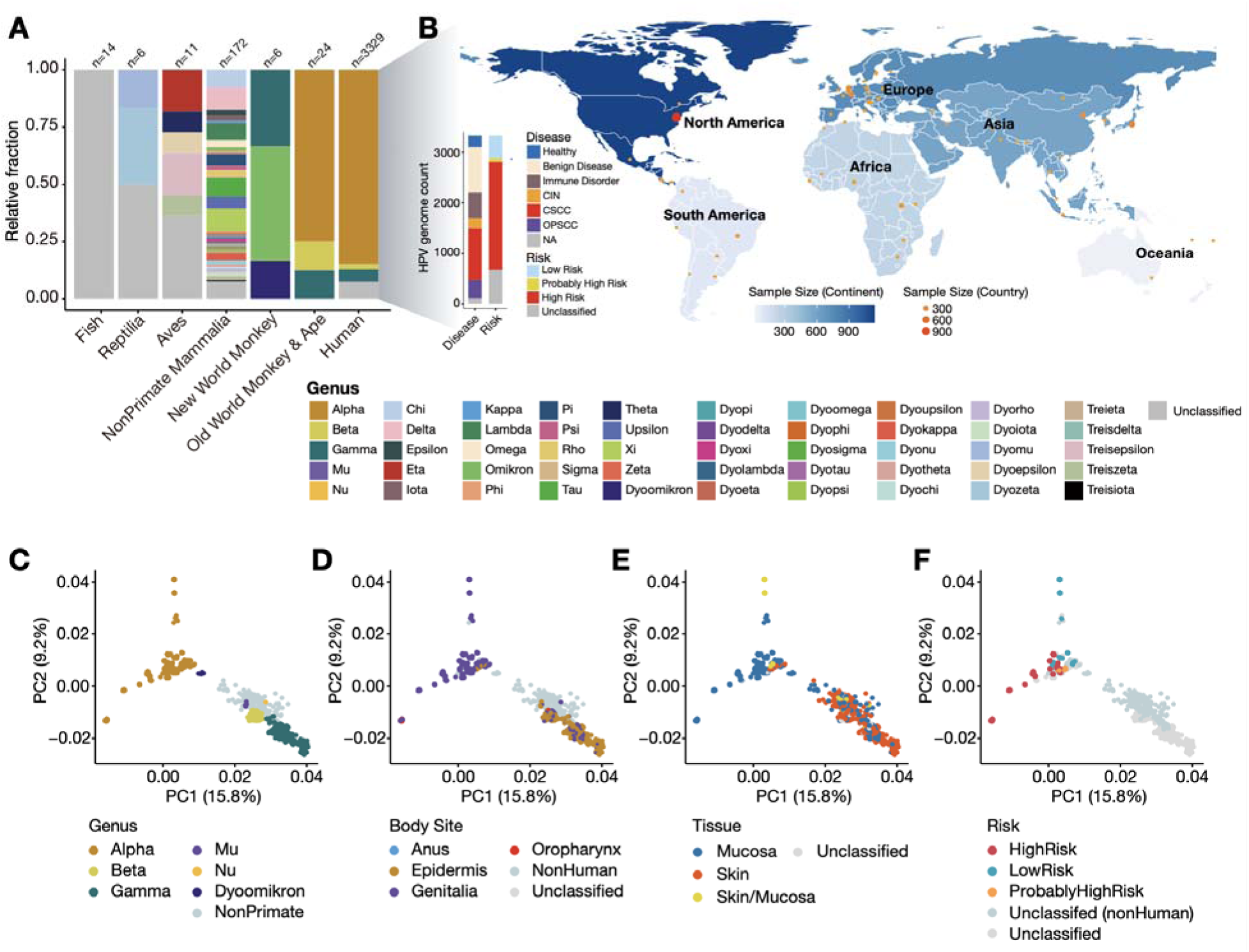
The global sampling and genomic diversity of the 3562 PV genomes. (A) The host and genus distribution of the 3562 PV genomes. (B) The geographical, disease type, and risk classification of the 3329 HPV genomes. (C–F) The principal component analysis of the global PV genome diversity with regard to PVs’ genus classification (C), anatomic body site origin (D), anatomic tissue site origin (E), and pathogenic risk classification (F). The detailed metadata of the plotted information is provided in Supplementary Table 1.

These 3562 PV genomes show rich variation in their genome content, genome size (4.7 kb–10.4 kb) and GC% (35.1%–59.2%). Regarding genome content, the *E1*, *E2*, *L1*, and *L2* genes are universally conserved and constitute the core gene set of PVs, whereas the other genes exhibited variable presence/absence patterns (e.g., the loss of E4, E1^E4, E8^E2 in bird PVs) (Supplementary Figure 1). As for genome sizes, fish PVs have clearly smaller genome sizes, largely due to their general depletion of the *E5*, *E6*, and *E7* genes (Supplementary Figure 2). Within the 3329 HPVs, significantly larger genome sizes were noticed for Alpha-PVs both before and after accounting for the phylogenetic relatedness, which are driven by expanded noncoding regions, especially the URR region that regulates viral transcriptional activity (Supplementary Figure 3). The extreme genome sizes of two HPV outliers are caused by structural variants (segmental deletion for HPV16_K2821 and segmental duplication for HPV11_GUMC_AJ_Lung), with the latter one experimentally verified in the original study [19] (Supplementary Figure 4).

We constructed all-against-all whole genome alignment for 3548 PV genomes from birds and terrestrial vertebrates and used the HPV type 16 (HPV16) as the reference to chart global genomic variation landscape of PVs with extracted genomic variants. The fish PVs were excluded for this analysis due to their large genomic divergence from other PV genomes. Principal component analysis (PCA) shows clear genomic differentiation between Alpha-PVs and PVs from other genera (Figure 1C). Such bipartition also holds in terms of anatomic body sites and tissue origins but not so much regarding host specificities, disease status, or geography (Figure 1D, E; Supplementary Figure 5). A clear distinction between high-risk and low-risk HPVs was further captured in our PCA analysis (Figure 1F). These observation collectively suggest substantial evolutionary divergence have taken place during the rise of Alpha-PVs a well as those high-risk HPVs at the genomic level, which likely opened opportunities for their niche expansion and pathogenesis.

### Time-resolved phylogeny of PV evolution

Our phylogenetic analysis on 3548 bird and terrestrial vertebrate PV genomes revealed a well-resolved phylogeny with 36 pure and 33 mixed clades (Supplementary Figure 6). Downsampled based on this phylogeny, we further generated a representative subset containing 600 PV genomes (including all 219 animal PVs and 381 representative HPVs covering all HPV lineages) to reduce redundancy in some heavily sequenced HPV types and built the corresponding downsampled PV tree (Supplementary Table 2). The 3548-genome-based and 600-genome-based PV trees show highly consistent topology, with PVs designated as the same type (e.g., HPV16) always form the same clade, while PVs designated with the same genus (e.g., Alpha, Beta, etc.) predominantly clustered together as well (Supplementary Figure 7). With the 600-PV-based tree, we evaluated the phylogenetic congruency between genome tree with gene trees derived from individual gene (e.g., *E1*, *E2*, *L1*, *L2*, etc.) or paired gene combinations (e.g., *E1-E2*, *E1-L1*, etc.) (Supplementary Figure 8, 9; Supplementary Table 3). The phylogenetic distances between the genome tree and gene tree negatively correlates with the sizes of the corresponding genes or gene pairs, with the gene pair E1-L1 showing highest consistency with the genome tree. When only considering single genes, E1-based gene tree better recaptured the genome-level phylogenetic signals compared with the more widely used L1-based gene tree.

Next we performed Bayesian molecular dating analysis on the 600-PV-based downsampled dataset to establish the evolutionary time frame of PVs (Figure 2; Supplementary Table 4). We estimated the divergence between the Alpha- and Beta/Gamma-super clades occurred at 96.7 million years ago (MYA). Given that Alpha-PVs are mostly mucosa-related whereas Beta/Gamma-PVs are predominantly skin-related, one can conjecture that such novel niche adaptation to mucosa is likely an important driver underlying such early divergence. The separation between the Beta- and Gamma-super-clades occurred more recently at 80.7 MYA. Interestingly, two bat PVs (min_sch_PV1 from the little bent-wing bat and ept_ser_PV2 from the serotine bat) branched off immediately before the Beta-Gamma divergence, hinting for the potential role of bats in mediating the early divergence of PVs. We noticed that the non-primate to primate host switch occurred within these three super clades roughly around the same time at 69.5-75.1 MYA, indicating a scenario of concordant non-primate-to-primate infection after the intra-host early divergence in non-primate mammals. Considering that the divergence time between human and the most closely related non-primate mammal flying lemur is 70.5-86.2 MYA, the primate PV infection likely occurred vertically along with the rise of primates. Within the primate PVs, the Alpha-PVs originated around 44.9 MYA, in sync with the separation of apes and old-world monkeys from new-world monkeys.

**Figure 2.**
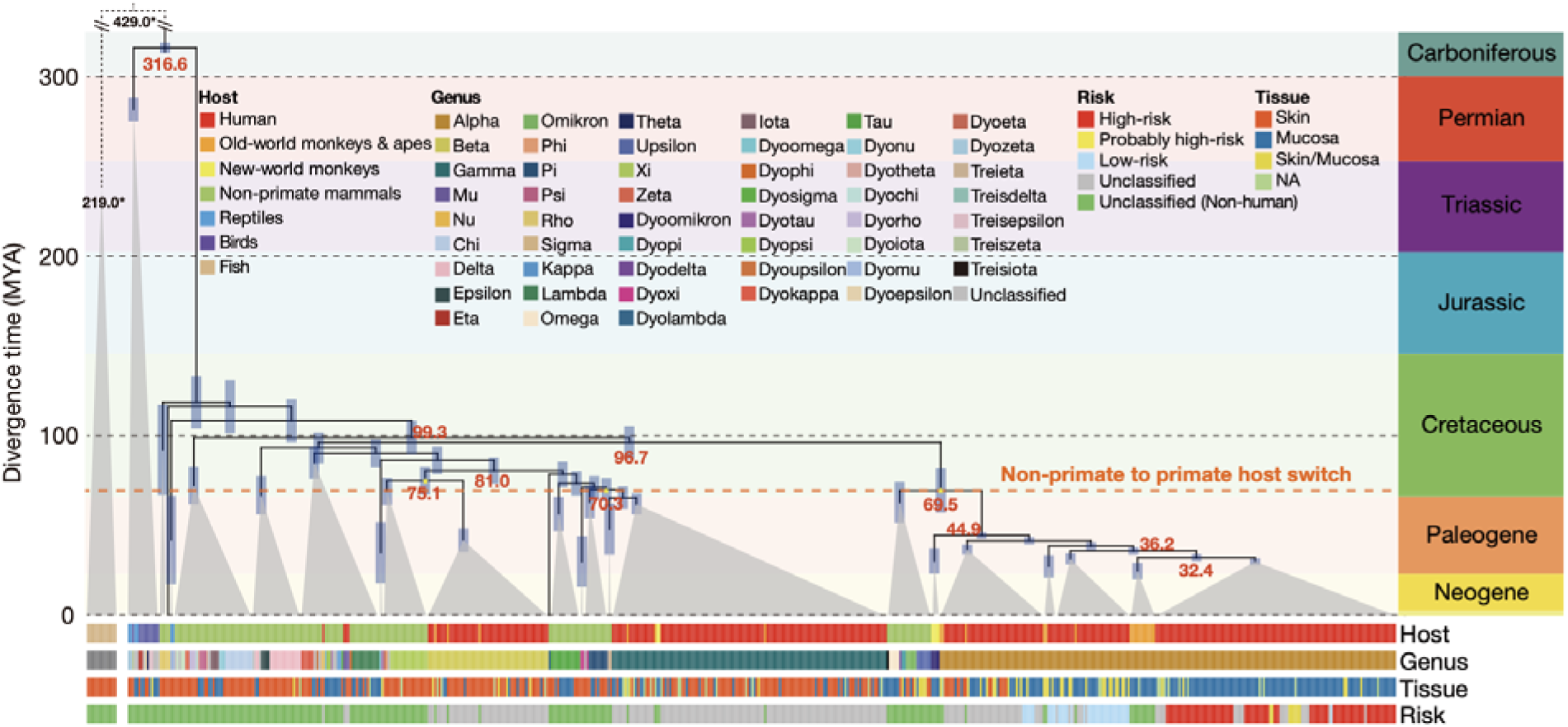
Evolutionary time frame of PV genome divergence and evolution. Calibrated time-resolved evolutionary tree for 600 representative PV genomes, with the branch lengths proportional to divergence time in million years ago (MYA). Key divergence time points regarding the early divergence of PVs were denoted in red, with the red dash line further highlighted concordant host shifts from non-primate mammals to primates for PVs from the Alpha, Beta, and Gamma super clades. The shades in each node represent 95% highest posterior densit (HPD) interval. The time tree of fish PVs was further annexed by referencing to their host divergence time (denote with asterisk). The tracks at the bottom show the phylogenetic distribution of PVs regarding their host species group, genus classification, tissue origin, and risk classification.

While PVs generally followed a virus-host co-divergence model, occasional instances of virus-host phylogenetic incongruency have also been reported [13,20]. Our analysis recaptured these observations, hinting for sporadic horizontal host switches. For example, some HPVs such as HPV1, HPV63, and HPV41 directly clustered with non-primate mammalian PVs from beaver and porcupine, while PVs from boa and python are deeply embedded in mammalian PVs rather than clustered with PVs from reptiles and birds. We employed a co-phylogenetic statistical framework to systematically assess such virus-host phylogenetic incongruence and to infer optimal evolutionary scenarios for reconciliation (Figure 3; Supplementary Table 5, 6). Our results revealed a consistently high level of duplication/loss turnover in PV genome evolution regardless of event-specific cost parameter settings (Figure 3A). We detected frequent horizontal host switch events involved with bats (13.6%), rodents/rabbits (18.2%), and primates (31.8%) (Figure 3B). For example, we found bats are strong candidates as the original source of HPV41 and its relative in porcupine, which are further nested within a bat-specific PV subclade that contains eight PVs from five bat species, hinting that bats should be the original source. Such a refined view on both when and how these host switch events occurred provides a new foundation for future investigation with enhanced PV sampling in these taxa.

**Figure 3.**
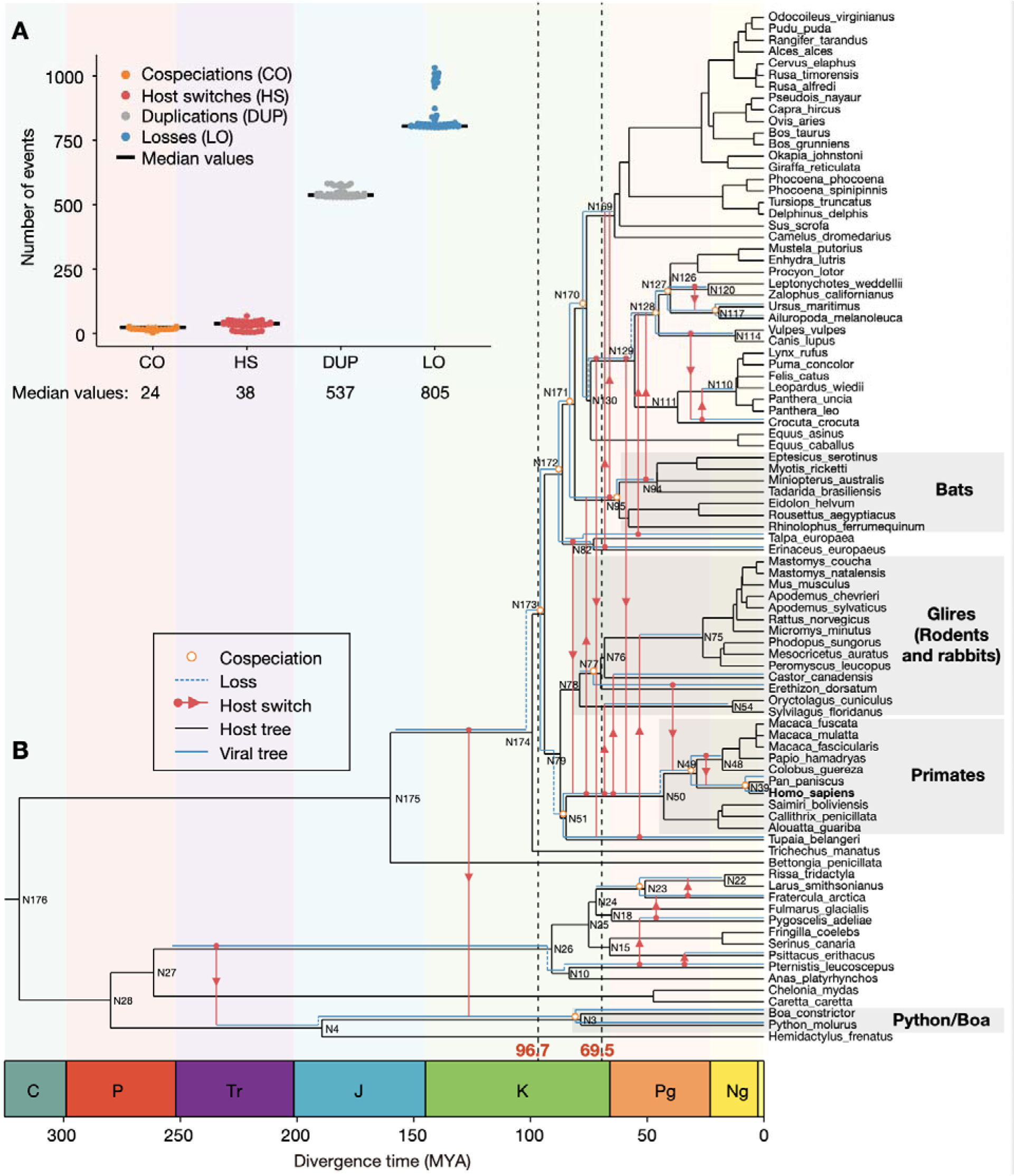
Phylogenetically reconciled evolutionary events during PV evolution. (A) The number of different types of evolutionary events inferred under a range of event-specific cost parameter settings. CO: cospeciation, HS: host switch, DUP: duplication; LO: loss. (B) Horizontal host switch events revealed by reconciliation between the virus and host phylogenetic trees. The phylogenetic node IDs associated with these host switch events were indicated, which were referred to in Supplementary Table 6. Geologically defined time zones are further depicted underneath. C: Carboniferous; P: Permian; Tr: Triassic; J: Jurassic; K: Cretaceous; Pg: Paleogene; Ng: Neogene. Key divergence time point such as 96.7 MYA for Alpha vs. Beta/Gamma PVs and 69.5 MYA for non-primate to primate host switch were also indicated.

### Selection driven mutations underlying the diversification among Alpha-, Beta-, and Gamma-PVs

Given the observed early divergence among Alpha-, Beta-, and Gamma-PVs, we set out to identify key mutations that define their genomic diversification. Following the phylogenetic tree, we screened for genomic variants that got progressively enriched in the Beta- and Gamma-super-clades as well as those got progressively depleted in the Alpha-super-clade (because the HPV16 reference genome used in our variant calling procedure belongs to the Alpha-super-clade). We postulated that these variant sites, whether in their derived or reference states, should represent the genomic diversification of the Alpha-, Beta-, and Gamma-PVs. In this way, we identified 29, 54, and 15 lineage-defining variants for the Alpha-, Beta-, and Gamma-super-clades respectively, along with another 11 variants shared by both Beta- and Gamma-super-clades (Figure 4A; Supplementary Table 7). Evaluated by mixed effects model of evolution (MEME) [21], we found 15 of these super-clade-specific genomic variants (corresponding to 12 amino acid substations in L1 and E1) under episodic diversifying selection (MEME test, *P*<0.1) (Figure 4B; Supplementary Figure 10). The fact that those with strong diversifying selection signals (i.e., MEME test *P*<0.05) are either Alpha-specific (n=5) or Beta-and-Gamma-specific (n=2) highlights the role of such site-specific selection in driving the early divergence of PVs (i.e., Alpha vs. Beta/Gamma). By using the bird and reptile PVs (clade M1) as the outgroup, we further determined the directions of these selection-driven mutations in evolution (Figure 4B; Supplementary Figure 10). Notably, the *E1* gene was strongly selected during the rise of Alpha-PVs, as all but one selection driven mutation occurred in *E1*, especially its encoded N-terminal regulatory domain (Figure 4C). Given that a major difference between Alpa PVs and Beta/Gamma PVs lies in their infected tissue type (mucosa vs. cutaneous epithelium), it is very possible that these N-terminal mutations contribute to such tissue-specific adaptation. The remaining selection-driven mutation that defines the Alpha-PV lineage locates in the *L1* gene, with its particular mutational site (L1: p.430 T->K) covered by an experimentally validated epitope (L1: p.427–445) [22,23] (Figure 4D). Future studies on these selection-driven lineage-specific mutations are expected to fully elucidate their functional and evolutionary impacts, especially in the context of Alpha-PV evolution.

**Figure 4.**
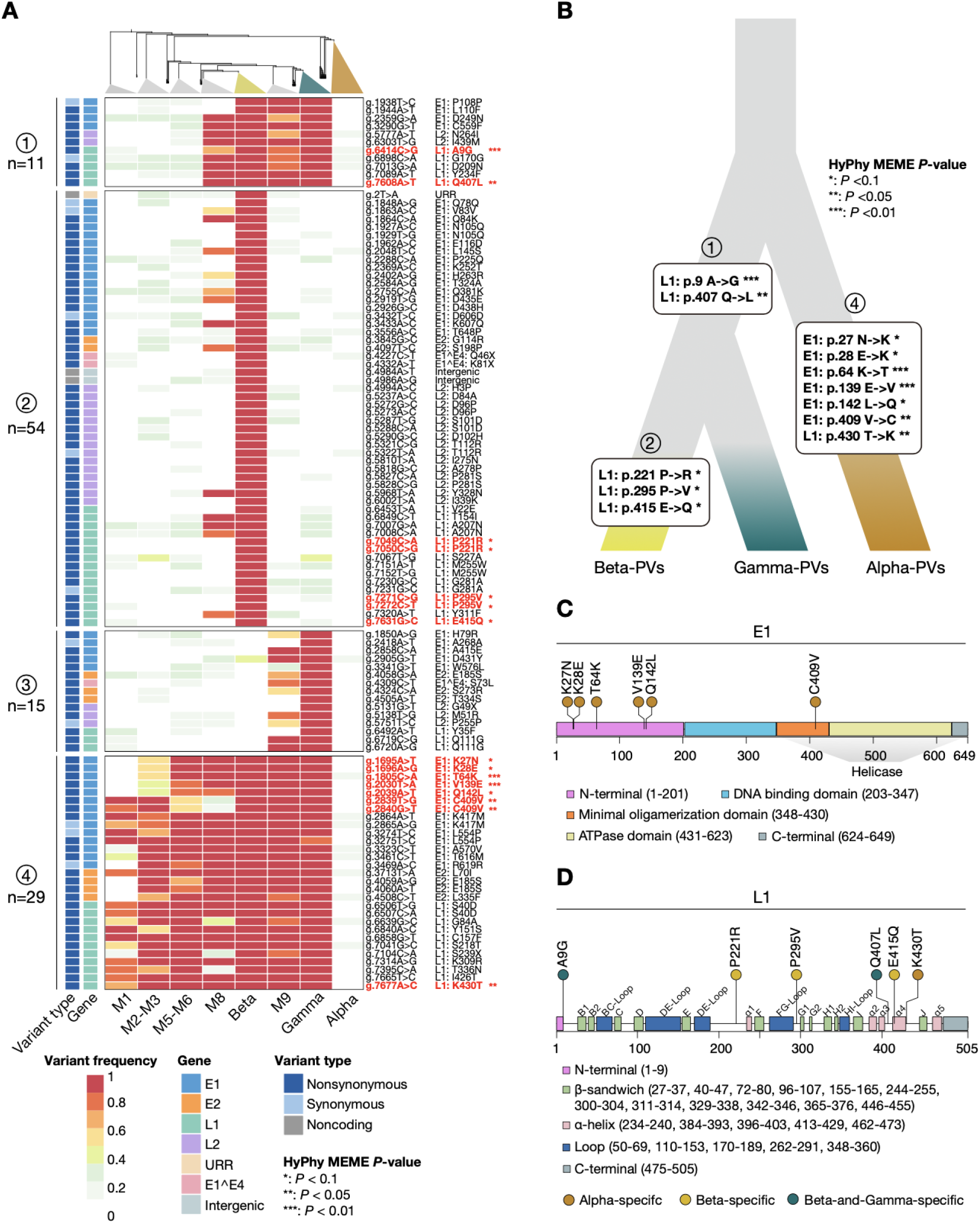
Key mutations defining the early divergence of the Alpha-, Beta-, and Gamma-PVs. (A) The prevalence heatmap of identified key mutations got progressively enriched in the Beta-and-Gamma-super-clades (⍰), Beta- (⍰), and Gamma-super-clades (⍰), or progressively depleted in the Alpha-super-clades (⍰). The genome and protein coordinates of these mutations were listed on the right, with the red fonts indicating mutations under significant diversifying selection (\**P*<0.1, \*\**P*<0.05, \*\*\**P*<0.01). (B) The phylogenetic distribution and mutational directions of key mutations under diversifying selection that drove the super-clade-specific diversification.

### Risk-defining HPV variant sites separating low-risk and high-risk HPVs

Our pangenome analysis reveals a highly correlated phylogenetic structure of HPVs regarding their pathogenicity classifications (Figure 5A, B). All “high-risk” and “probably high-risk” HPVs tightly clustered into a single derived lineage within the Alpha-super-clade, suggesting a lineage-specific origin of high pathogenicity. Referring to our molecular dating analysis, we estimated these high-risk HPVs arose around 32.4-36.2 MYA. The two most common high-risk HPVs, HPV16 and HPV18, got diversified at 1.4 MYA and 0.6 MYA respectively, suggesting a scenario of ancient high-risk HPV diversification (both across and within different HPV types) in archaic hominins, before the rise of modern human [24].

**Figure 5.**
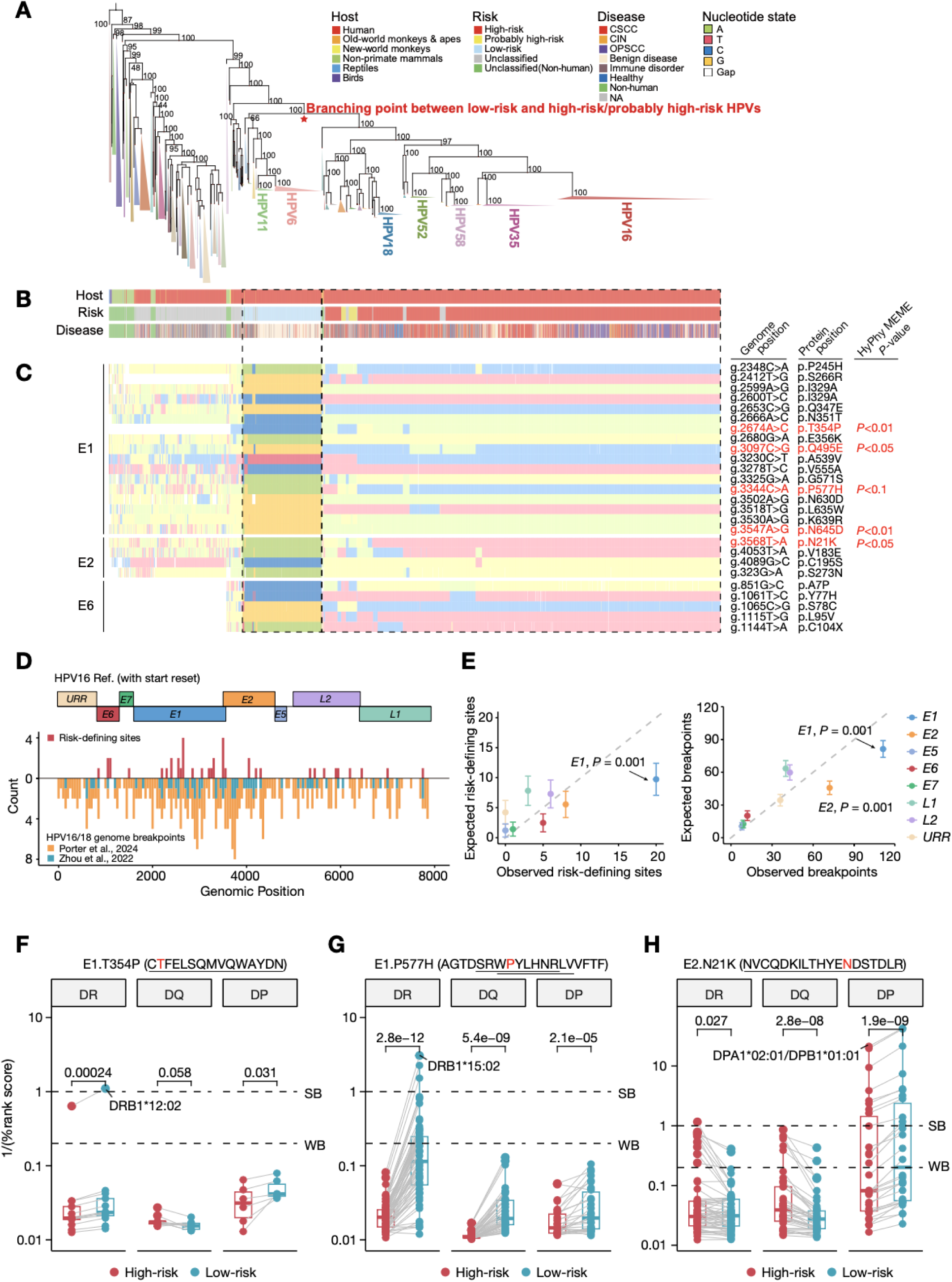
Risk-defining variant sites with contrasting genotypes between low-risk and high-risk HPVs. (A) The phylogenetic tree of 3548 PVs. (B) The metadata track for the 3548 PVs regarding their species origin, pathogenic risk, and associated diseases. (C) The nucleotide state of risk-defining variant sites in *E1* and *E2*. Sites highlighted in red are subject to significant diversifying selection evaluated by HyPhy MEME. (D) The physical distribution of risk-defining variant sites and high-risk HPV genome breakpoints. (E) Permutation test of gene-specific enrichment of risk-defining sites (left) and high-risk HPV genome breakpoints (right). (F-H) The MHC-II binding affinity comparison of epitopes between high-risk and low-risk epitopes in E1 (F, G) and E2 (H). The amino acid changes in epitope peptides (underlined sequences) were indicated above, while the MHC-II binding affinity [measured by 1/(%rank_score)] for the top 3 MHC-II alleles from nine geographical regions were plotted below. The MHC-II allele with the highest binding affinity were highlighted. The *P*-value of Wilcoxon signed-rank test (paired, two-sided) is given. Dashed lines represent the thresholds for strong binding (SB, 1%) and weak binding (WB, 5%).

To identify key mutations associated with the rise of high-risk PVs, we first evaluated the branch-specific selection scheme around the branching point between low-risk and high-risk HPVs, which was tested by adaptive branch-site random effects likelihood (aBSREL) on a gene by gene basis [25]. We found significant episodic diversifying selection (aBSREL test, *P*<0.05) in *E1* and *E2* genes immediately after this branching event for the high-risk HPV lineages as well as in E2 and E6 along the following branch (Supplementary Figure 11). Since *E1* and *E2* genes are essential for HPVs’ viral genome replication and transcription, and *E6* is a classic oncogene, it is no surprising that their coding sequence was positively selected along with the rise of high-risk HPVs. At site-specific level, we found a total of 40 variant sites (34 nonsynonymous and 6 synonymous) showing contrasting genotypes between low-risk and high-risk HPVs (Figure 5C; Supplementary Figure 12; Supplementary Table 8). We refer these 40 variant sites as risk-defining sites given their genotype segregation between high-risk and low-risk HPVs.

More than half of these risk-defining sites locate in either *E1* or *E2* genes, with their genomic positions in close adjacency to the hotspots of high-risk HPV genome breakpoints introduced during host-integration [26,27] (Figure 5D). Our permutation tests also suggest a significant enrichment of both risk-defining sites and HPV genome breakpoints in *E1* gene (*P*=0.001) (Figure 5E). These observations collectively suggest an intrinsic association between our identified risk-defining sites and HPV genome breakpoints, potentially shaped by the microhomology-mediated DNA repair process accompanied with high-risk HPVs’ host genome integration. In addition, natural selection also contributed to the observed contrasting genotypes between low-risk and high-risk HPVs as we detected site-specific diversifying selection at five nonsynonymous risk-defining variant sites (MEME test, *P*<0.1). These five sites locate in E1’s helicase domain and E2’s transactivation domain, both of which are critical for HPVs’ viral replication activity. At the structural level, mutations at these sites also notably altered on E1’ and E2’ original intramolecular interactions (Supplementary Figure 13). Aside from such direct functional impacts on E1 and E2, specific genotypes selected at these sites could also contribute HPVs’ pathogenicity by promoting immune evasion. We found three (two in *E1* and one in *E2*) of these five selection-driven risk-defining sites overlapped with six experimentally verified immune epitopes[28–31] (Supplementary Table 9), for which epitopes carrying the high-risk variant forms exhibit generally lower affinity to major histocompatibility complex class II (MHC-II) molecules compared with those carrying the low-risk variant forms (Figure 5F-H; Supplementary Table 9, 10). This finding is concordant with previous reports from genome-wide association studies (GWAS) suggesting that MHC-II genes are strongly associated with the subjectivity of cervical and oropharyngeal cancers, two most notable HPV-associated cancer types[32–34]. Therefore, the selected genotypes at these risk-defining variant sites in high-risk HPVs likely confer them with stronger potential for escaping from the host immune surveillance.

### Clinical implications of HPV variants in pathogenic risk and prognosis prediction

Conventionally, the pathogenetic risk of an HPV isolate is scored based on its specific subtype (e.g., high risk for HPV16 and low risk for HPV6). Given the strong segregating patterns between low-risk and high-risk HPVs in our identified risk-defining variant sites, it could be possible to predict high-risk vs. low-risk HPVs by just knowing the genotypes of these sites. Here we systematically evaluated their prediction value in pathogenic risk prediction with four independent cohorts, which include: 1) 194 HPVs genomes annotated as “complete” in Genbank, 2) 5871 HPV genomes annotated as “partial” in Genbank, 3) 131 HPV genomes sequenced from HPV16-positive cervical cancer (cervical squamous cell carcinoma, CSCC) patients examined in this study, and 4) 63 HPV genomes sequenced from HPV16-positive oropharyngeal cancer (oropharyngeal squamous cell carcinoma, OPSCC) patients examined in this study (Supplementary Table 11-14). We verified the HPV genomes that we assembled with cohort 3 and 4 are indeed HPV16 genomes based on a phylogenetic analysis covering all of them as well as the 600 PV set (Supplementary Figure 14). For each of our identified 34 nonsynonymous risk-defining variant sites, we examined how well could our identified representing genotypes for high-risk or probably high-risk HPVs in our pangenome analysis be recovered by the high-risk or probably high-risk HPVs included in the corresponding validation cohort (Figure 6A, B; Supplementary Table 15-18). Here we used a set of widely used classification metrics such as precision, recall, and F_1_ score for this benchmark. For cohort 1 and 2, both low-risk and high-risk HPVs are included, therefore all these three metrics can be readily calculated. Regardless which metric was used, we found 27 (for cohort 1) and 24 (for cohort 2) of the 34 signature sites exhibit satisfying performance scores >=0.9, indicating well-balanced sensitivity and specificity in HPV risk classification even just based on just a single variant site. For cohort 3 and 4, since only high-risk HPV genomes (HPV16) were included, we calculated the recall value for each risk-defining variant site. For almost all sites (32/34 for CSCC and 33/34 for OPSCC), we got perfect recall values of 1.0 with these two cohorts, which again highlights the application value of our identified risk-defining variant sites in predicting HPVs’ pathogenic risk.

**Figure 6.**
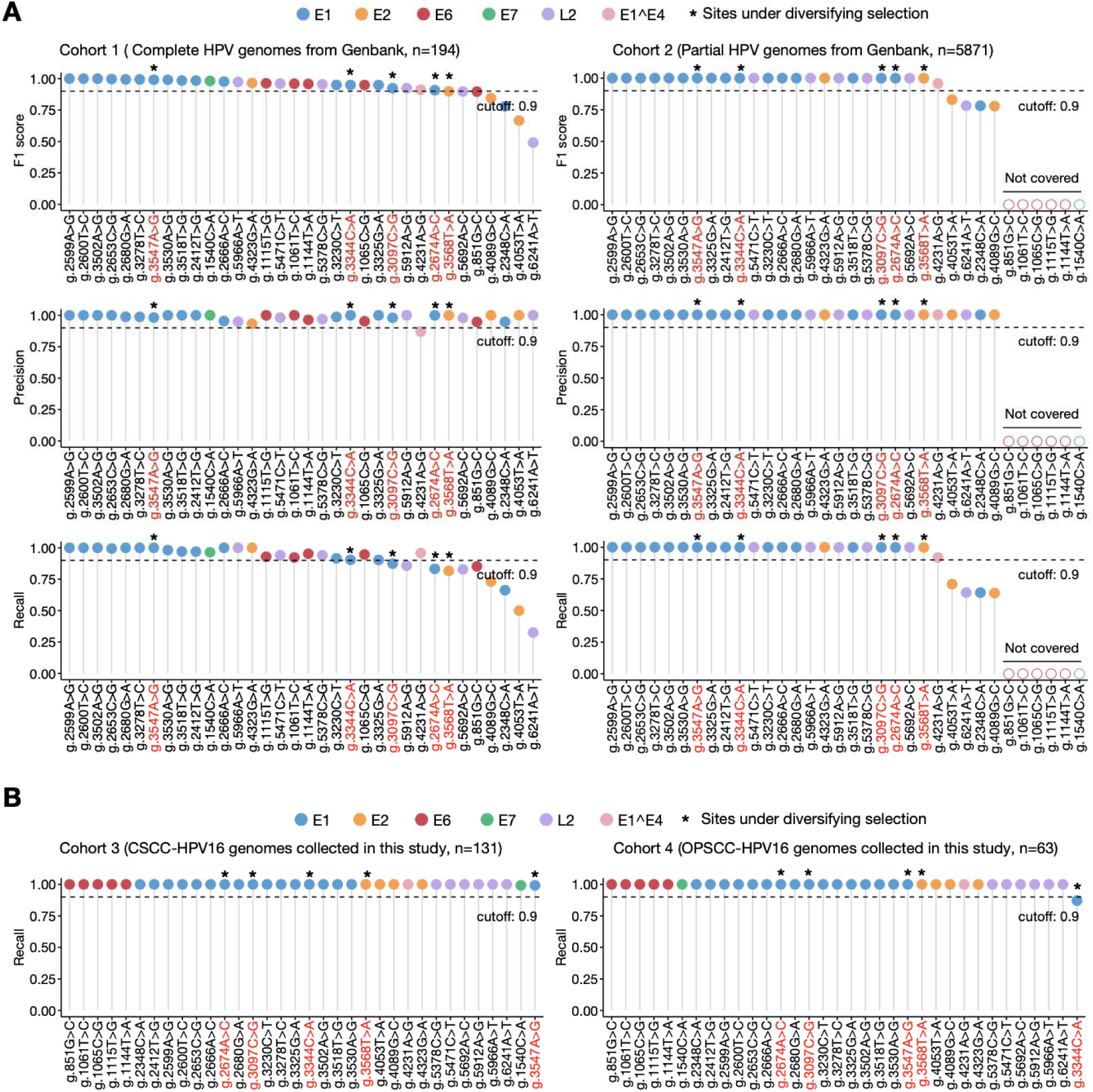
Performance validation of HPV risk-defining variant sites for pathogenic risk prediction. (A) The prediction performance of each individual risk-associated signature site in identifying HPVs with high-risk or probably high-risk pathogenicity in cohorts retrieved from Genbank (left: cohort 1; right: cohort 2). (B) The prediction performance of each individual risk-associated signature site in identifying HPVs with high-risk or probably high-risk pathogenicity in cohorts collected in this study (left: cohort 3; right: cohort 4). The five selection-drive sites are further highlighted in red color as well as with the asterisk notation. The genomic coordinates indicated are based on the start-reset-version of the HPV16 reference genome.

In addition to risk prediction for HPVs, the identification of biomarkers that predict patient survival is also of importance in clinical practice. For HPV-associated cancers, the positivity of high-risk HPVs (e.g., HPV16) has been used as a primary prognostic predictor in this regard[35–37]. Until recently, much less is known regarding the prognostic value of specific variants in high-risk HPV genomes on their associated cancer patients[38]. With the HPV16-positive CSCC (n=131) and OPSCC (n=63) patient samples collected in this study (cohort 3 and cohort 4 aforementioned; mean follow-up time: 5.39 year for CSCC and 5.08 year for OPSCC), we identified four HPV16 variants (three for CSCC and one for OPSCC) with significant impacts on patient overall survival (OS) after the false discovery rate (FDR) correction (Figure 7A-D, Supplementary Table 19-20). The three CSCC-prognostic variants (two in L2 and one in L1) show strong linkage to each other, with univariate hazard ratios ranging from 10.34 to 15.46 (Supplementary Figure 15). As for OPSCC, our identified one prognostic variant locates in E2 with a univariate hazard ratio of 22.71. Multivariate Cox regression analysis on these four prognostic HPV16 variants suggested added predictive accuracy in addition to traditional prognostic factors such as age and tumor stage (Figure 7E). Two (L2 p.306 and E2 p.198) of these four prognostic variants are nonsynonymous, and they both show notable changes in intramolecular interactions at the structural level (Figure 7F, G; Supplementary Figure 16). Taken together, although with relatively small cohort size, our analysis provides a proof of concept that some HPV variants could have significant impact on the survival of HPV-associated cancer patients. Future studies with larger human cohorts from distinct ethnic groups are expected to reveal more such variants that could be used collectively towards a better prognosis prediction.

**Figure 7.**
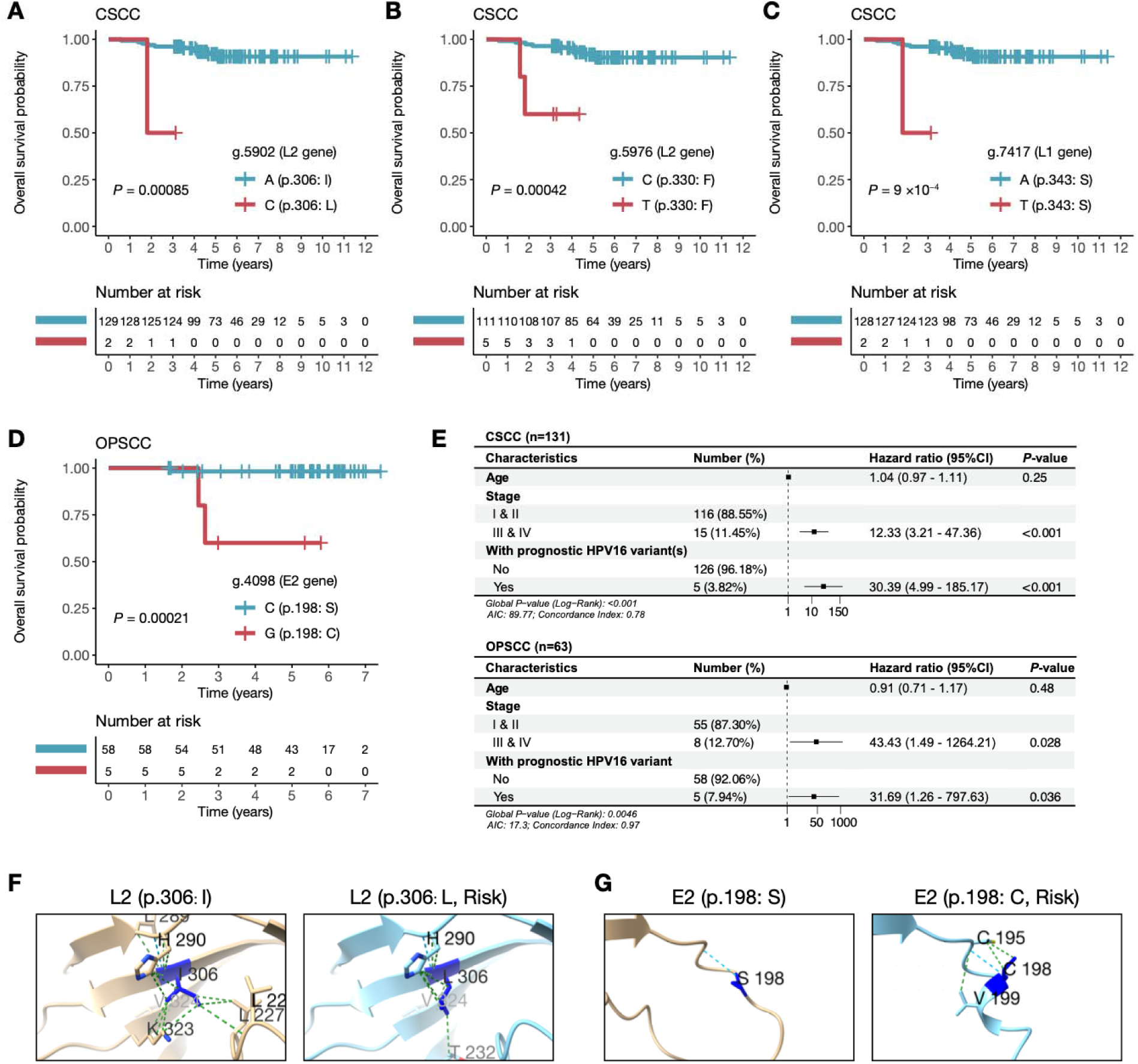
Discovery of prognosis-associated HPV16 variants. (A–D) The overall survival (OS) of patients stratified by individual prognostic HPV16 variants identified in our CSCC (A–C) and OPSCC (D) cohorts. (E) Prognostic factors for overall survival (OS) by multivariate analysis. (F,G) The structural view of the two nonsynonymous prognostic variants identified in *L2* (F) and *E2* (G) and their impacts on intramolecular interactions

## Discussion

PVs are a group of ancient viruses that have evolved over millions of years and infected a wide range of animal hosts, including human. With such long-term evolution, it is theoretically expected that the pathogenicity of viruses tend to mitigate to reach a state of chronic inapparent infection, especially for the slowly evolved double stranded DNA viruses such as PVs[39]. Indeed, this seems to be the case for PVs evolved in most animal lineages. However, specific types of human PVs (i.e., HPVs) such as HPV16 and HPV18 have been proven to be highly carcinogenic, imposing a global public health threat to human society. Therefore, it is important to study the genetic and molecular basis of such pathogenicity evolution. The rapid progress of sequencing technologies and their increasing adoption in HPV-positive patients and animal hosts in recent years vastly expanded the genomic resources of PVs available[18,40], enabling a comprehensive comparative study between low-risk and high-risk HPVs through the lens of genome evolution.

In this study, we systematically collected and curated 3562 human and animal PV genome sequences to generate a super-pangenome collection of PVs reflecting their global diversity. Using comparative genomics and molecular evolution frameworks, we established a pangenomic and time-resolved view for PVs’ genome evolution. We identified clear genomic differentiation between Alpha-PVs and PVs from other genera is correlated with their unique body parts and tissue specificity. Especially, we found genomic sites under Alpha-PVs’ lineage-specific selection predominantly fell in the N-terminal regulatory domain of the E1 protein, highlighting the importance of this domain in promoting Alpha-PVs’ niche adaptation regarding both body parts and tissue specificity. Consistent with our finding, previous studies found this N-terminal regulatory domain of E1 from anogenital HPV but not with cutaneous HPV or animal PVs can interact with the cellular WD repeat protein p80 (WDR48), therefore linking HPVs’ N-terminal domain with their niche specificity[41,42]. Future functional study on our identified selection-driven alterations in E1’s N-terminal domain is expected to provide more mechanistic insights into this aspect.

This leads to the next question, what genomic changes in evolution confer the high pathogenicity for some of Alpha-PVs during their association with human. We found 40 genomic variant sites (34 nonsynonymous and 6 synonymous ones) in the HPV genome showing strong segregating pattern in terms of their genotypes between low-risk and high-risk/probably high-risk HPVs, which further agreed well with the phylogenetic bifurcation between these two groups of HPVs. We consider these variant sites as risk-defining ones and argue that they are highly informative in telling high-risk and low-risk HPVs apart even at the single site level, without knowing any information from the rest of the HPV genome. These sites have important application value in both primary (vaccination) and secondary (screening) prevention against high-risk HPVs. As a proof of concept, we brought in four independent validation cohorts containing >6300 HPV genomes in total and found our identified risk-defining variant sites well-performed in differentiate low-risk versus high-risk HPVs in all four cohorts. Interestingly, more than half of these 40 risk-defining sites fall in *E1* and *E2* gene rather than the *E6* and *E7* genes that are directly oncogenic, which might seem counterintuitive at the first glance. Given that E1 and E2 are essential for the replication and regulatory control of viral life cycle [43–46], we think our observed pattern might reflect a deeper evolutionary divergence underlying HPVs’ pathogenicity evolution. Mechanistically, we found both mutational and selective forces likely contributed to the segregating genotype pattern at these sites. For example, we detected physical association between these sites and high-risk HPV breakpoints, suggesting microhomology-mediated DNA repair as a potential mutagenic source in shaping this pattern. Meanwhile, we found at least some of these risk-defining variant sites are under significant diversifying selection, reinforcing the genotype segregation between high-risk and low-risk HPVs. For these sites, we found the specific genotypes carried by high-risk HPVs can contribute to a higher potential of immune evasion, which explains why they are associated with high-risk pathogenicity. Collectively, these findings highlight how the virus-host interaction is shaping HPVs’ genome and pathogenicity evolution.

Given the strong association between high-risk HPV infection and malicious human cancers such as cervical and oropharyngeal cancers, discovery and application of clinically relevant HPV genomic variants in high-risk HPVs could have great value in the prevention and treatment of HPV-associated cancers. In this study, we provided proof-of-concept examples demonstrating how these variants can be used in predicting HPVs’ pathogenic risks as well as the prognosis of HPV-associated cancers. While the clinical cohorts collected in our current study are still relatively limited, it is nevertheless encouraging to see how some of our identified key HPV variants can be informative in real clinical settings. We expect future studies with larger and broader HPV-associated clinical cohorts from more diverse ethnical groups can further enrich our understanding on the biological and clinical significance of these key HPV variants in the age of precision medicine.

Finally, most existing PV studies, including our current one, are anthropocentric, meaning that our sampling and knowledge of PVs are heavily biased towards human. In comparison, our knowledge of the genetic and ecological diversity of animal PVs are still rather limited, which prevents a high-resolution view into their deep evolutionary history. Recently, several large national and international consortia have been formed to sequence and characterize genomes of a wide variety of eukaryotic life on earth[47,48]. Aside from contemporary animals, researchers have also began to look for trace of PV infection in ancient animals [49]. Taken together, it can be expected that our knowledge on the virome of the life on earth could be extensively expanded in both taxonomically and historical sampling. When such data becomes available, the community will be able to assemble a more balanced and complete view on PVs regarding its biology and evolution.

## Materials and Methods

### PV genome collection and curation for publicly available data

A total of 2969 complete HPV genomes (quality control criteria: assembly gap <12.5% of the total length) were downloaded from the Genbank database hosted by the National Center for Biotechnology Information (NCBI) via the NCBI EFetch utility[50] (v16.2) with the query term of “human papillomavirus complete genome” (last accessed by 2023.06.01). Another 360 HPVs genomes (quality control criteria: assembly gap <12.5% of the total length) were retrieved by contacting the authors of a recently published study[38]. In addition, 233 animal PVs (including 30 non-human primate PVs, 172 non-primate mammal PVs, 6 reptile PVs, 11 bird PVs, and 14 fish PVs) were retrieved from the PaVE database[18,40] (last accessed on 2023.06.01). The metadata regarding PVs’ host species origin, PVs’ anatomic body parts and tissue origin, patients’ geography origin, PV’s taxonomic classification, PV’s risk classification, patients’ disease type classification, etc. was further retrieved and curated by referencing back to the original studies that sequenced these PV genomes. Given the different starting site choice and annotation quality of these PV genomes, we further reset the starting site of each assembly as the first base after the L1 stop codon and performed gene annotation with PuMA[51] (v1.2.2). The software gffread[52] (v0.9.12) was further used to extract the coding sequences (CDSs) from each annotated gene accordingly.

### 3548-PV-genome-based whole-genome alignment and phylogenetic tree building

All curated PV genomes were processed with windowmasker[53] (v1.0.0) for soft-masking. MAFFT[54] (v7.310; options: --auto --preservecase) was used to generate a preliminary whole-genome alignment for the soft-masked 3548 curated human and animal PV genomes (including 2968 HPVs, 30 non-human primate PVs, 172 non-primate mammal PVs, 6 reptile PVs, 11 bird PVs). The 14 fish PV genomes were excluded due to their substantially large sequence divergence with other PVs, which tends to compromise the alignment quality. The resulting alignment was processed by ClipKIT[55] (v1.3.0; options: -m gappy -g 0.9) before being fed to IQ-TREE[56] (v2.0.6; options: --seed 20230417 -m MFP -B 1000 --bnni -alrt 1000 --safe) to generate the guide tree, which was further used to generate the second-round whole-genome alignment with Cactus[57] (v2.2.0) by using the start-reset HPV16 (hom_sap_HPV16; NC_001526.4 with start-reset) as the reference genome. The Cactus derived hal-formatted alignment were converted to the maf format by the hal2maf tool from the HAL package[58] (v2.2.0; options: -- onlyOrthologs --noDupes --maxRefGap 2000 --refGenome hom_sap_HPV16), which was further converted to the fasta format. The resulting fasta-formatted genome alignment was clipped by ClipKIT[55] (v1.3.0; option: -m gappy) and manually inspected before providing to IQ-TREE[56] (v2.0.6; options: -- seed 20230421 -m MFP -B 1000 --bnni -alrt 1000 --safe) for the final tree building. The resulting tree was rerooted by the nw_reroot tool from the Newick Utilities[59] (v1.6.0) with PVs from birds and reptiles as outgroups. The final phylogenetic tree was visualized using FigTree (v1.4.4; http://tree.bio.ed.ac.uk/software/figtree/) and ggtree[60] (v3.6.2).

### 600-PV-genome-based whole-genome alignment, phylogenetic tree building, and molecular dating analyses

Based on the phylogenetic structure and host species distribution of 3548-PV-genome-based phylogenetic tree, we further generated a down-sampled dataset with 600 representative PV genomes. Such PV genome down-sampling was performed within each type-specific HPV clade for retaining the overall phylogenetic characteristics while reducing the computational burden for downstream analysis. Whole-genome alignment and tree building were further carried out based on this down-sampled 600 PV genomes following the same procedures that we used for the 3548-PV-genome-based analysis. According to previously estimated PV evolutionary rates[61], we used a Bayesian Markov Chain Monte Carlo (MCMC) based framework implemented by BEAST2[62] (v2.5.2) for molecular dating analysis. Following the suggestions from the similar analyses that have done previously[12,63,64], we used a combination of Yule tree prior and relaxed uncorrelated lognormal distribution (UCLD) of the molecular clock model for our BEAST analysis. Regarding the calibration nodes, we introduced four calibration point along the phylogenetic tree based on a collective consideration of the general codivergence histories between PVs and their hosts, as well as the PV-specific the early pre-divergence of the Alpha, Beta, and Gamma superclades in non-primate mammal hosts: (1) the separation between the reptile PVs (represented by hem_fre_PV1/2) and Aves PVs (represented by che_myd_PV1) at 280.0 (274.9–286.8) MYA, (2) the split between reptile/bird PVs (e.g., hem_fre_PV1/2, lar_smi_PV1, pyg_ade_PV1/2) and mammalian PVs (e.g., rou_aeg_PV1, equ_cab_PV1) at 319.0 (95% CI of 316.0–322.4) MYA, (3) the split between new-world monkey PVs (e.g., sai_sci_PV1/2/3, alo_gua_PV1) and old-world monkey PVs (e.g., mac_mul_PV3, mac_fus_PV2) at 43.0 (95% CI of 40.0–44.2) MYA, and (4) the split between the lineage-specific human (e.g., hom_sap_HPV13) and chimpanzee PV (e.g., pan_pan_PV1) at 6.2 (6.0–6.5) MYA. These divergence time were retrieved from the TimeTree database[65] (v5; https://timetree.org/). Two independent MCMC runs were launched for 200000000 MCMC steps, with 10% burn-in and a subsampling frequency of 20000 generations. A minimal effective sample size (ESS) of 200 was used for evaluating the MCMC chain convergence, which were inspected in Tracer (v1.7.1; http://tree.bio.ed.ac.uk/software/tracer/). TreeAnnotator (part of with BEAST2 package) was used to generate the consensus tree with 10% burn-in. The resulting consensus tree was visualized with FigTree (v1.4.4; http://tree.bio.ed.ac.uk/software/figtree/).

### Genome size comparison accounting for phylogenetic relatedness

To account for phylogenetic non-independence in the PV genome size comparisons across different genera, we fitted a Bayesian Phylogenetic Linear Mixed Model (PGLMM) using the *R* package brms[66] (v2.22.0). First, the phylogenetic tree was converted into a variance-covariance matrix using the vcv.phylo function in the *R* package ape[67] (v5.8-1). The PGLMM was specified with the following formula: genome_size ∼ genus + (1|gr(PV_name, cov = cov_matrix)), in which genome_size is the response variable and genus is the fixed effect predictor. The model assumed a Gaussian error distribution.

Four independent Markov chains were run, each with 4000 iterations in addition to 1000 iterations of burn-in. After confirming model convergence (Rhat ≈ 1.00), we extracted the phylogenetically corrected marginal mean genome size for each genus using the *R* package emmeans[68] (v1.11.2) and carried out the pairwise comparisons among all compared genera. A statistically significant difference was considered if the 95% credible interval of the posterior distribution for the difference did not overlap with the zero line.

### Variant calling and annotation

Single nucleotide variant calling was identified from the final alignment of 3548 PV genomes using SNP-sites[69] (v2.5.1; option: -mv). The raw calling set were filtered using the following criteria: (1) variants with a missing rate of >80%, (2) variants with a missing rate of >10% and a minor allele frequency (MAF) of <0.01, with the latter filtering step carried out by PLINK[70] (v1.9; options: --geno 0.1 --maf 0.01 -- allow-no-sex --make-bed --const-fid). We defined the variant passing the first filter as the “base variant set” and those passing both filters as the “core variant set”. Next we used Ensembl VEP (v109.3) to annotate and predict the functional impacts of these variants based on a minorly edited HPV16 genome annotation GFF file, in which 1) genomic features labeled as “polyA_signal_sequence”, “TATA_box” and “URR” were relabeled as ncRNA_genes (so that variants fall in these region will all be denoted as “non_coding_transcript_exon_variant” by Ensembl VEP) and 2) genes only with “CDS” annotations but not the associated “mRNA” and “exon” annotations were fixed. After the Ensembl VEP prediction, we curated the results with the following criteria: 1) For variants annotated as “non_coding_transcript_exon_variant”, we reclassified them as “regulatory_variant”; 2) For variants annotated as “upstream_gene_variant” and “downstream_gene_variant”, only the annotations correspond to the closest gene were considered; 3) For variants located between coding genes, we labeled them as “intergenic_variant”; 4) For variants associated with multiple genes (e.g., overlapping genes), we denoted the variant based on the gene with higher functional impact.

### Principal component analysis

We performed principal component analysis (PCA) on the “core variant set” using the smartpca function from the EIGENSOFT package[71] (v8.0.0). First, the VCF-format file was converted to PLINK-format files using PLINK[70] (v1.9; options: --make-bed --allow-no-sex --double-id). The generated PLINK files was then converted to input files of smartpca using the convert function from EIGENSOFT (options: outputformat: EIGENSTRAT). Finally, the resulting files were used to perform PCA analysis using smartpca (options: lsqproject: YES numoutlieriter: 0).

### PV tree versus host tree reconciliation

To infer the horizontal host switch events based on the associations between PVs and their hosts, we used Jane (v4) [72] to compare the 600-PV-genome-based tree against the time-resolved phylogenetic tree of their hosts (retrieved from the TimeTree database[65]). The host and the 600-PV trees were converted into a Jane-compatible timed tree in nexus format using the Python script available on GitHub[73] (https://github.com/andersonbrito/openData/tree/master/brito_2020_reconciliation). For the Jane analysis, a 5-million-year “time zone” was used to order different evolutionary events. Also, a set of pre-defined relative cost parameters for different evolutionary schemes need to be provided, for which we tried multiple combinations of relative cost values (e.g., co-speciation [0], duplication [0–3], loss [0–3], and horizontal host switch [0–3]) to have a good sampling for the parameter space. In this way, a total of 64 solutions for cophylogenetic reconstruction were obtained. The host switch event that occurred in ≥80% of the solutions was defined as a high-confidence event.

### Lineage- and risk-defining variant enrichment analysis

We performed the lineage- and risk-defining variant enrichment analysis using hypergeometric test on the “base variant set” defined previously with the Benjamini-Hochberg false discovery rate (FDR) correction. For lineage-defining variants, we identified them using the following criteria: 1) FDR-corrected *P*-value <0.01 in the focal vs. background hypergeometric test, 2) variant allele frequency ≥90% in the focal lineage, 3) variant allele frequency <20% in all non-focal lineages. Note that because the HPV16 genome that we used as the reference belongs to the Alpha-PVs, the lineage-defining variants for Alpha-PVs were identified as those got progressively depleted in this clade. For risk-defining variants, we scanned for variants that are enriched in either low-risk HPV groups or high-risk HPV groups. For the former, we used the criteria of 1) FDR-corrected *P*-value <0.01 in the low-risk HPV group vs. probably high-risk/high-risk HPV group hypergeometric test, 2) variant allele frequency ≥95% in the low-risk HPV group, 3) variant allele frequency <10% in the high-risk HPV group. For the latter, we used the criteria: the criteria of 1) FDR-corrected *P*-value <0.01 in the high-risk and probably high-risk HPV group vs. low-risk HPV group hypergeometric test, 2) variant allele frequency ≥95% in the high-risk and probably high-risk HPV group, 3) variant allele frequency <10% in the low-risk HPV group. Additionally, we further require the major allele genotype frequency in the high-risk and probably high-risk HPV group to be >=80%. Finally, for the case where multiple variant sites hitting the same codon, we used get_MNV (v1.0.0; https://github.com/PathoGenOmics-Lab/get_MNV) to identify and relabel the corresponding amino acid changes accordingly.

### Branch- and site-specific selection analysis

We used aBSREL[25] from the HyPhy package[74] (v2.5.60) to detect episodic diversifying selection in a branch-specific manner along the 3548-PV-genome-based phylogenetic tree and its associated alignment for the Alpha-super-clade and its subtrees, with the following options: --kill-zero-lengths No -- branches Foreground. In parallel, we also used the mixed effects model of evolution (MEME) [21] implemented in HyPhy to detect specific sites underlying diversifying selection across different HPV genes (e.g., *E1*, *E2*, *E5*, *E6*, *E7*, *L1*, and *L2*).

### HPV genome breakpoints analysis

To understand the association between risk-defining sites and HPV-host genome integration breakpoints. We used HPV16/18 genome breakpoints characterized in two studies using long-read sequencing and mapped these genomic positions to the start-reset HPV16 genome using liftOver[75] (v482). Next, we used the 50-bp non-overlapping window to profile the distribution of both risk-defining sites and HPV16/18 breakpoints across the HPV16 genome. To determine whether the observed sites were randomly distributed, we used the shuffle command from the BEDTools package[76] (v2.30.0) to generate 1,000 sets of randomly shuffled sites across the HPV genome, based on which an empirical *P*- value of region-specific enrichment was further calculated for the *URR*, *E1*, *E2*, *E5*, *E6*, *E7*, *L1*, and *L2* regions.

### Epitopes affinity prediction

We first retrieved information of experimentally verified HPV16 immune epitopes from IEDB[77] (https://www.iedb.org/), with which we further extract those peptide epitopes covering our identified risk-defining variant sites for epitope binding affinity prediction. Meanwhile, we retrieved the human MHC allele frequencies of nine geographical regions (Europe, North Africa, North America, South America, South Asia, Sub-Saharan Africa, East Asia, Western Asia, and Oceania) from the AFND database[78] using HLAfreq [79] (v0.0.4). For each geographical region, the top 3 alleles of each MHC gene were used for epitope binding affinity prediction. Given the previously reported strong association between MHC-II and HPV-associated cancers, we employed NetMHCIIpan[80] (v4.3) to predict MHC-II binding affinity to the epitope peptides that carrying either the wildtype or mutated form of our query sites with default parameters. Wilcoxon signed-rank test (paired, two-sided) was used to compare the overall epitope-to- MHC-affinity between the wildtype and mutated forms.

### Protein structure prediction and visualization

To evaluate the structural impact of the mutation, the original and mutated amino acid sequences of our interested HPV16 protein were independently submitted to the AlphaFold3 server[81] (https://alphafoldserver.com/) for structure prediction. The resulting crystallographic information files (CIFs) were then imported into ChimeraX [82] (v1.8) for structural comparison and visualization. Interactions involving the mutated residue and surrounding amino acids were analyzed using the “hbonds” and “contacts” commands in ChimeraX.

### Clinical sample collection

The study was approved by the ethics review board of the Sun Yat-sen University Cancer Center (SYSUCC) (Issue number: B2022-509-01). A total of 131 HPV16-positive CSCC and 63 HPV16-positive OPSCC patient samples were collected from the Tumor Biobank, Department of Gynecologic Oncology, and Department of Head and Neck Surgery of SYSUCC. All patient samples passed our quality control for DNA extraction, sequencing, and genome assembly (quality control criteria: assembly gap<12.5% of the total length). For each clinical sample, frozen tissues derived from surgically treated patients with follow-up data were included. The 8th edition American Joint Commission on Cancer (AJCC) staging manual[76] was referred to for staging the CSCC and OPSCC patients. The CSCC patients are all females, with an average age of 54 years old (ranging from 29 to 80) and most of them being clinically classified as stage I-II (89%, 116/131). The OPSCC patients are mainly male (87%, 55/63) with an average age of 55 years old (ranging from 33 to 70), most of which being staged at I-II (87%, 55/63).

### Cohort validation of HPV risk-defining variant sites

The predictive power of the identified HPV risk-defining variant sites for pathogenic risk was benchmarked using F1 score, precision, and recall. These metrics were calculated with a custom Python script. The classification of an allele at a given risk-defining variant site as a “high-risk allele” was considered a positive prediction. For this analysis, true positives (TP) were defined as the number of high-risk alleles identified in HPV strains classified as high-risk/probably high-risk types. False positives (FP) were the number of high-risk alleles found in HPV strains classified as low-risk types. False negatives (FN) were the count of non-high-risk alleles (excluding gaps) within the high-risk/probably high-risk HPVs.

### HPV genome amplification and sequencing

All patient samples were tumor tissues freshly frozen in liquid nitrogen within 30 min of resection. Genomic DNA extraction was performed by using the FastPure Cell/Tissue DNA Isolation Mini Kit (Cat: DC102-01, Vazyme, China). To minimize the possibility of cross-contamination, no more than 10 samples were processed in the same batch. Based on the extracted genomic DNA, we used Touchdown PCR (TD-PCR) or Nested PCR to amplify the complete genome of HPV16 with six sets of previously published primers[77] (Supplementary Figure 17; Supplementary Table 21) to increase the specificity and sensitivity. Between TD-PCR and Nested PCR, TD-PCR was prioritized. Our TD-PCR mix includes 25 μL Green Taq Mix (Cat: P131, Vazyme, China), 2 μL of TD-PCR forward and reverse primer (referred as F1/F2/F3_Touchdown_F/R in Supplementary Figure 19; 10 μM), ∼80 ng genomic DNA and sterile DEPC-H2O (Cat: R0022, Beyotime, China), which adds up to a final volume of 50 μL. Our TD-PCR program is as follows: 95°C 3 min; touchdown phase (2 cycles: 95°C 15 sec, 65°C 15 sec, 72°C 3 min 10 sec; 2 cycles: 95°C 15 sec, 62°C 15 sec, 72°C 3 min 10 sec; 4 cycles: 95°C 15 sec, 59°C 15 sec, 72°C 3 min 10 sec); 30 cycles: 95°C 15 sec, 56°C 15 sec, 72°C 3 min 10 sec; 72°C 5 min. As for nested PCR, the first-round amplification was carried out by using 12.5 μL Green Taq Mix, 1 μL of round 1 forward and reverse primer (referred as F1/F2/F3_NestedRound1_F/R in Supplementary Figure 19; 10 μM), 40–80 ng genomic DNA and sterile DEPC-H_2_O in 25 μL reaction. The PCR product of the first-round PCR was diluted with 75 μL sterile DEPC-H_2_O and was used as a template for second-round amplification, which was performed in a 25-μL reaction mix containing 12.5 μL of Green Taq Mix, 1 μL of round 2 forward and reverse primer (referred as F1/F2/F3_NestedRound2_F/R; 10 μM), 5 μL template and 5 μL sterile DEPC-H_2_O. The same PCR cycling program was used for both rounds: 95 °C for 5 min, 20 cycles at 95 °C for 30 sec, 56 °C for 30 sec and 72 for 3 min 10 sec. Final extension was set at 72 for 5 min. For PCR product clean-up, we used the 1× VAHTS DNA Clean Beads (Cat: N411, Vazyme, China). The obtained HPV16 amplicons, containing three purified PCR amplicon fragments which were examined by electrophoresis and further quantified by Qubit 4 Fluorometer (Thermo Fisher, USA). For each sample, we mixed the three PCR amplicon fragments with a 1:1:1 mass ratio and sent for sequencing by BGI Genomics (China) using the BGI MGISEQ-2000 platform.

### HPV genome assembly

The raw sequencing reads were preprocessed using fastp[83] (v0.24.0) to remove adaptor and low-quality bases. The trimmed reads were aligned to a reference genome that contains both the host (chm13v2.0) and viral genome sequence (HPV16, NC_001526.4) using BWA[84] (v0.7.17; option: mem). Only the paired-end reads mapped to the viral genome were selected for the *de novo* genome assembly with SPAdes[85] (v4.2.0; option: --isolate). All viral reads were then scaffolded with RagTag[86] (v2.1.0). Gap filling was performed using Abyss-sealer[87] (v2.3.5, options: -b20G -k32 -k48 -k64 -k80 -k96 -k112 -k128). After gap filling, assembly polishing was performed using Pilon[88] (v1.24, options: --fix snps,indels,local). A finally quality check was performed based on the dotplot generated by mummerplot[89] (v3.5) against the HPV16 reference genome. Similarly, we reset the starting site of each assembly as the first base after the L1 stop codon and performed gene annotation with PuMA[51] (v1.2.2). For each patient-derived HPV16 genome assembly, we used MAFFT[54] (v7.310, options: --auto --preservecase) to align it to the start-reset HPV16 genome and used SNP-sites[69] (v2.5.1) to extract variant sites, which were further annotated by Ensembl VEP[90] (v109.3).

### Prognostic analysis

The prognostic analysis was carried out with the *R* package survival (v3.7.0) and further visualized with survminer (v0.5.0). The Cox proportional hazards model was used to evaluate the association between each patient-carrying HPV16 variant (minor allele frequency ≥1%) and overall survival. To estimate the risk of death from any cause, the only censoring event was at the time of last known follow-up, and the years since cancer diagnosis was used as the time variable. The Benjamini-Hochberg false discovery rate (FDR) was calculated based on the raw *P*-values for multiple test correction. An FDR-corrected *P* ≤ 0.05 (two-tailed) was considered statistically significant. Multivariable Cox regression analysis was used to explore the association between the bearing status of our identified prognostic HPV16 variant (i.e., with or without) and overall survival, further adjusted for widely used clinical prognostic factors such as age and stage. The forestploter package[91] (v1.1.3) was used for result visualization.

## Acknowledgments

We thank Dr. Dmitri Petrov for the insightful comments and feedback to this work. This work is supported by National Natural Science Foundation of China (32470663 to Jia-Xing Yue and 32000395 to Jing Li), Guangdong Provincial Pearl River Talents Program (2019QN01Y183 to Jia-Xing Yue and 2021QN02Y168 to Jing Li), Natural Science Foundation of Guangdong Province (2022A1515010717 to Jia-Xing Yue and 2022A1515011873 to Jing Li), and Young Talents Program of Sun Yat-sen University Cancer Center (YTP-SYSUCC-0042 to Jia-Xing Yue and YTP-SYSUCC-0040 to Jing Li). The funders have not played any role in the study design, data collection and analysis, decision to publish, or preparation of the manuscript.

## Author contributions

Huihui Li, Jingtao Chen, and Xinxin Zhang contributed equally. Jia-Xing Yue and Jing Li conceived the study. Huihui Li, Jingtao Chen, Xinxin Zhang, Ludong Yang, Lin Feng perform the data analysis. Xinxin Zhang, Jingtao Chen, Yan Shuai, Congying Chen performed the molecular experiment. Jihong Liu, Ming Song, Xin Zhang, Yi Ding contributed to the clinical sample collection. Jia-Xing Yue, Jing Li, and Huihui Li wrote the paper. All authors reviewed and contributed to the final version of the paper.

Correspondence to JL or JXY.

## Data availability statement

The curated genome assemblies, genome annotations, alignments, variant sites, raw and time-resolved phylogenetic trees of the 3562 publicly available PVs, as well as their associated metadata and bioinformatics scripts, altogether packed as Supporting Dataset 01-07, have been deposited at zenodo under the DOI of <u>10.5281/zenodo.10049935</u>. The sequencing reads of the HPV16 genomes isolated from the 131 CSCC and 63 OPSCC patients collected in this study have been deposited at the Genome Sequence Archive (GSA) (https://ngdc.cncb.ac.cn/gsa/) in National Genomics Data Center[92] under the accession number PRJCA020850 and the reads accession numbers <u>CRA013231</u> and <u>CRA027844</u>. The corresponding genome assemblies of these HPVs have been submitted to GenBase[93] (https://ngdc.cncb.ac.cn/genbase/) under the accession numbers C_AA111526.1 to C_AA111719.1. respectively.

