## Supplementary Figure 1-17 for "Super-pangenome analysis of 3562 human and animal papillomavirus isolates illuminates their genome and pathogenicity evolution"

**Supplementary Figures**

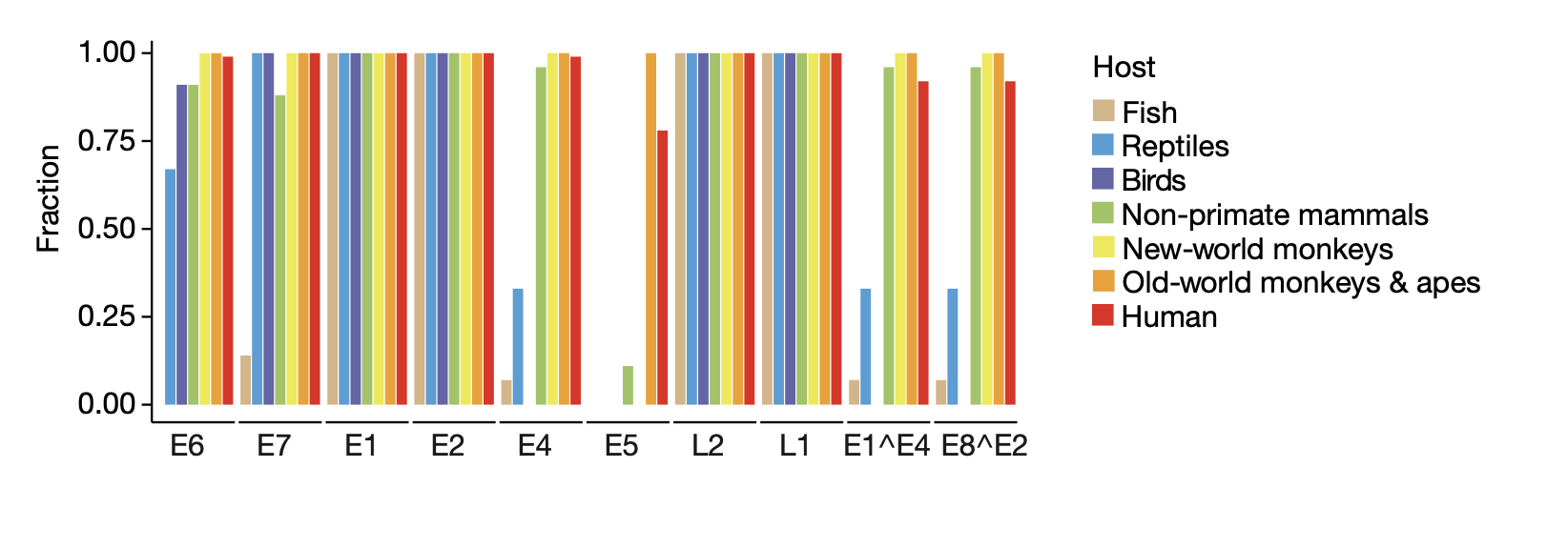

**Supplementary Figure 1 The genome content variation of PVs from different host taxonomic groups.**

The prevalence ratio of each PV gene in each host taxonomic group (indicated by different colors) is shown.

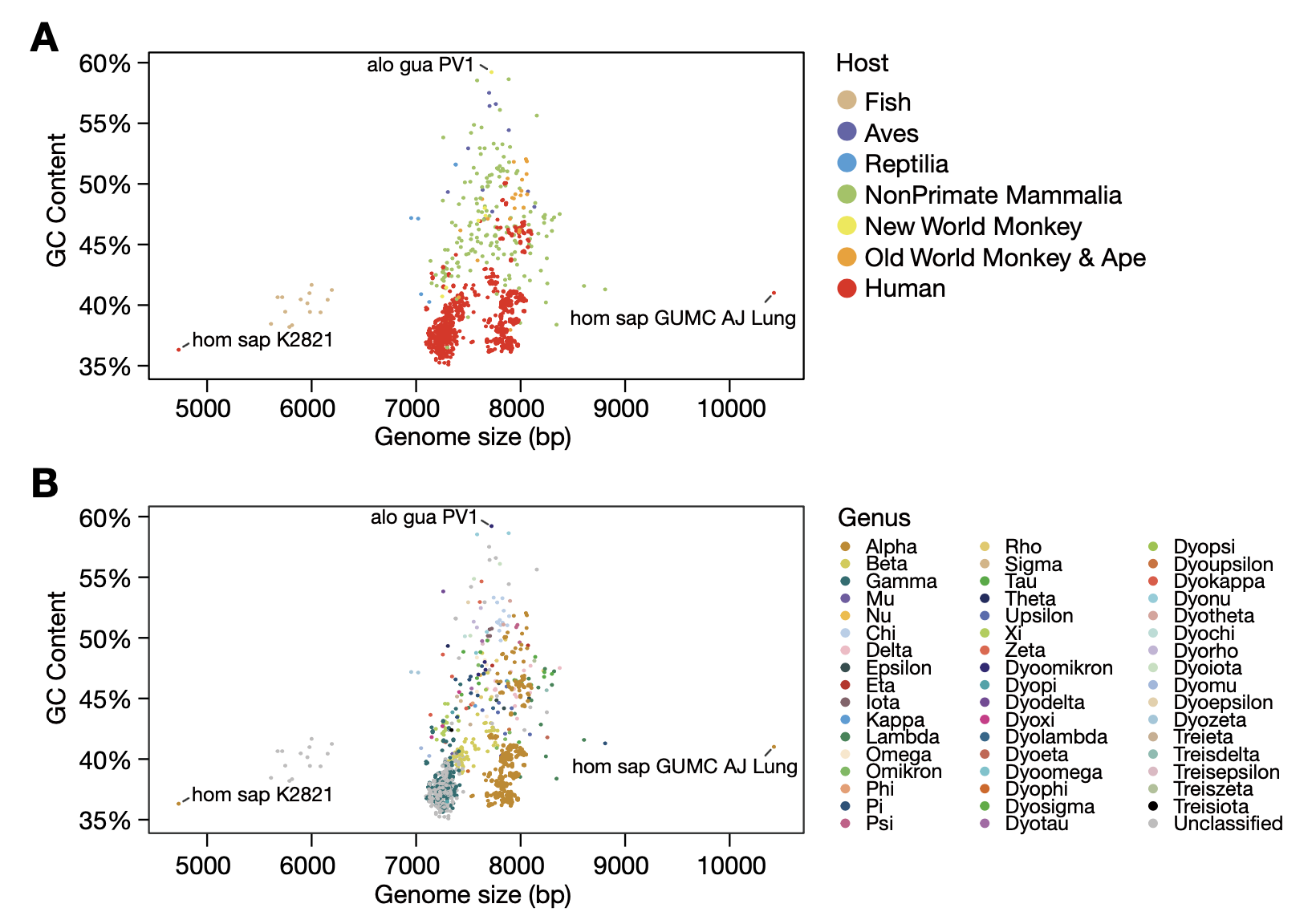

**Supplementary Figure 2. Genome size and GC content variation for the 3562 PV genomes.**

The genome size and GC content distribution of the 3562 PV genomes were plotted with regard to their host (A) and genus (B) classification.

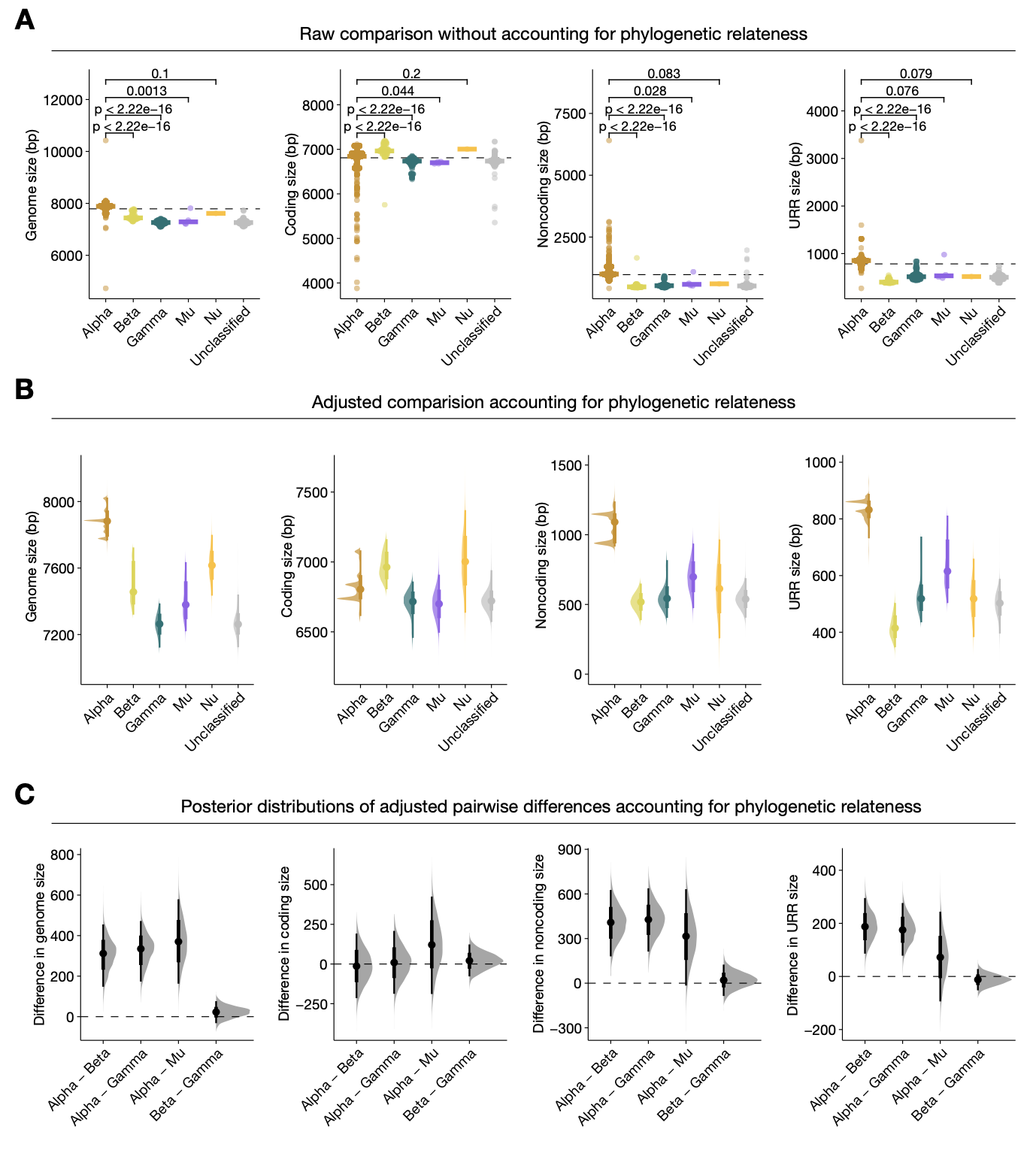

**Supplementary Figure 3. Size comparison of the whole genome, coding, and noncoding regions among 3329 HPVs regarding their designated genera.**

(A) Raw comparison based on two-sided Wilcox rank sum test. (B) Adjusted comparison accounting for phylogenetic relatedness using phylogenetic linear mixed model (PGLMM). (C) Posterior distribution of pairwise differences of PVs from different genera estimated by PGLMM.

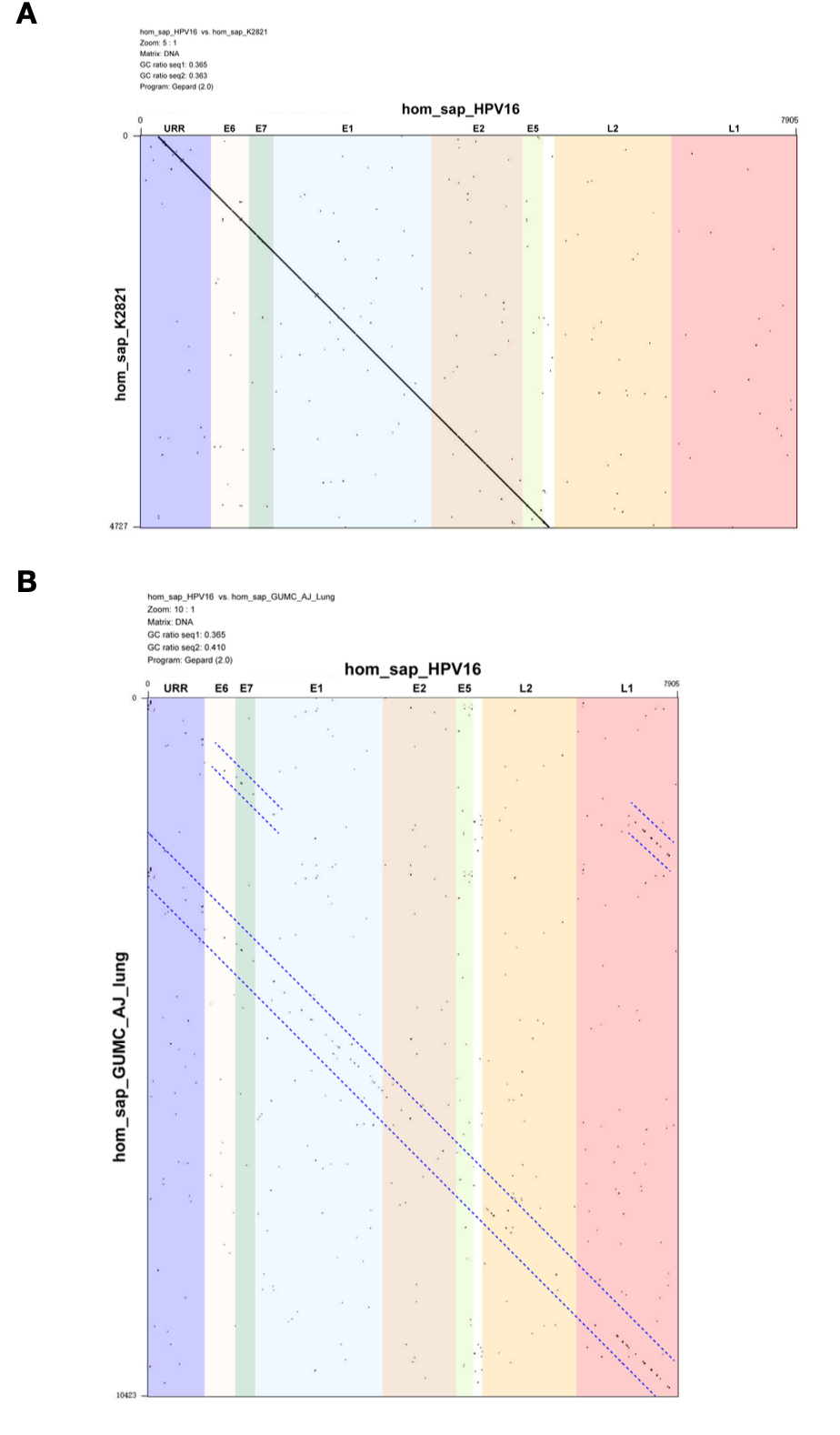

**Supplementary Figure 4. The structural variants in hom_sap_K2821 and hom_sap_GUMC_AJ_lung.**

The genome comparison dotplots for hom_sap_K2821 (A) and hom_sap_GUMC_AJ_lung (B) are shown against hom_sap_HPV16 respectively. The paths of the main colinear signal are highlighted by blue dashed lines.

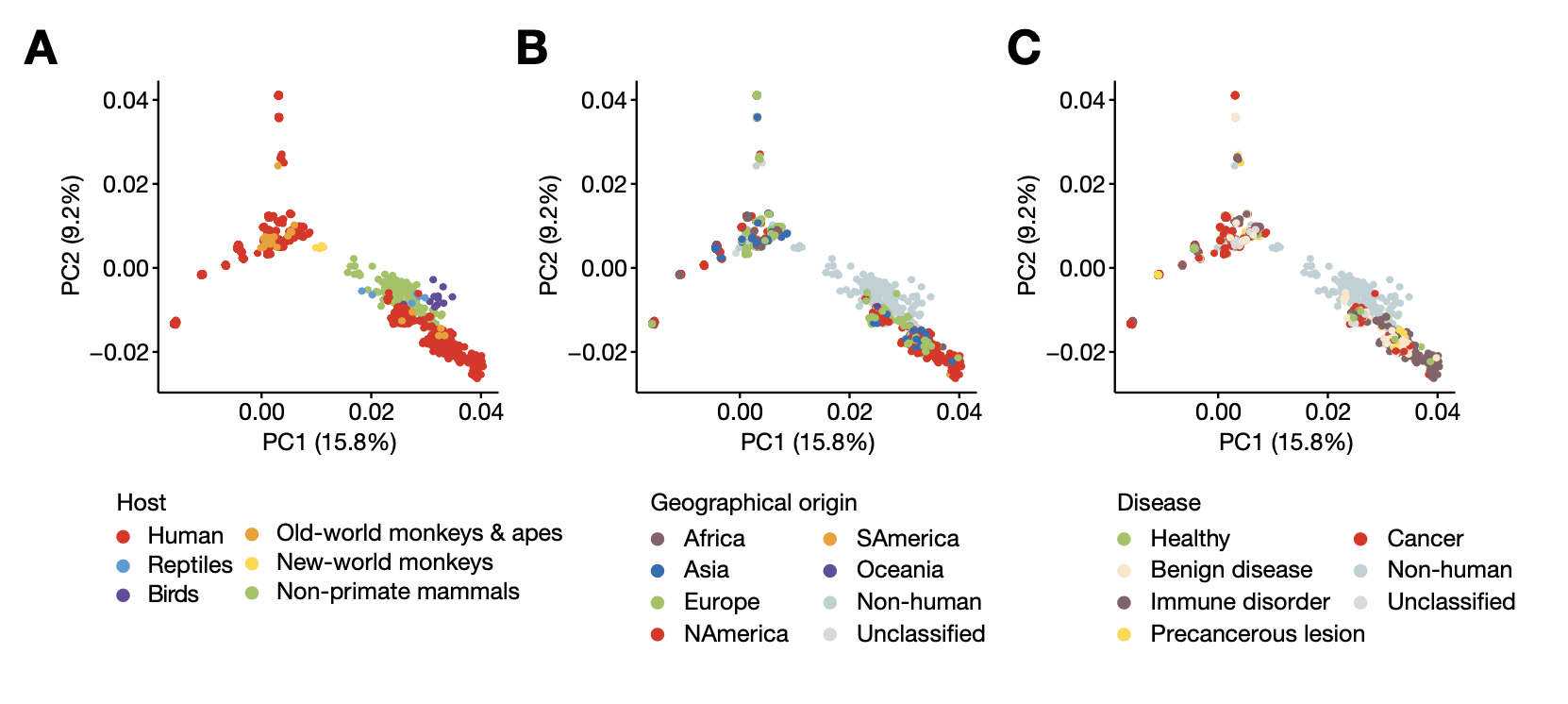

**Supplementary Figure 5. Principal component analysis using genomic variants extracted from the 3548 PV genomes.**

Data points were color-labeled by host species (A), geographical origin (B), and disease status (C) respectively.

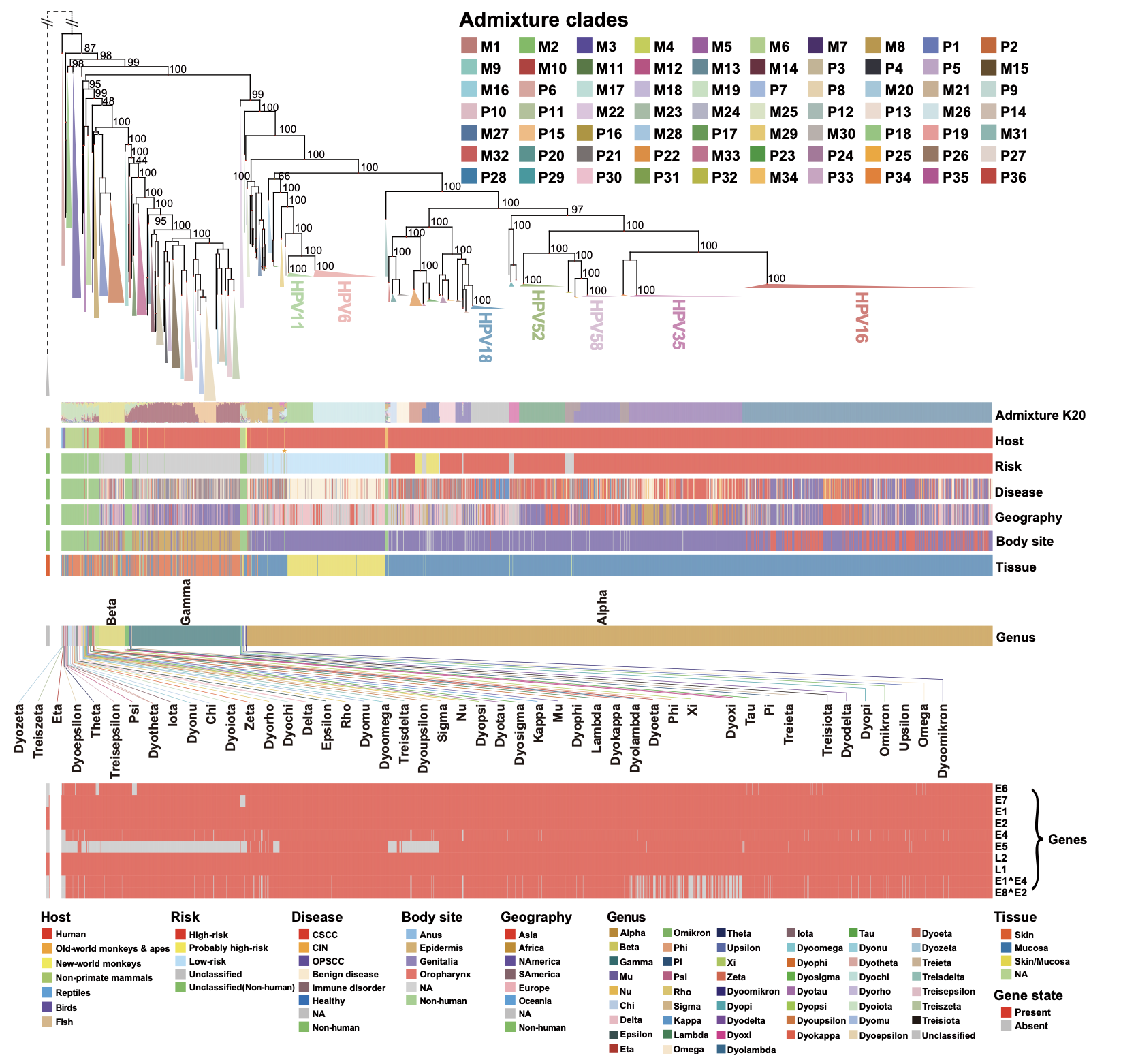

**Supplementary Figure 6. The phylogenetic tree of 3562 PVs.**

The maximum likelihood phylogenetic tree was built based on the whole-genome alignment of 3548 PVs and 14 fish PVs respectively, and the two trees were annexed together afterwards. The labeled node support values were calculated by approximate likelihood-ratio test (aLRT). Below the phylogenetic tree, individual tracks were displayed regarding host species groups, risk classification, disease types, geographical origins, anatomic body site origins, tissue origins, genus classification, and gene content (from above to bottom).

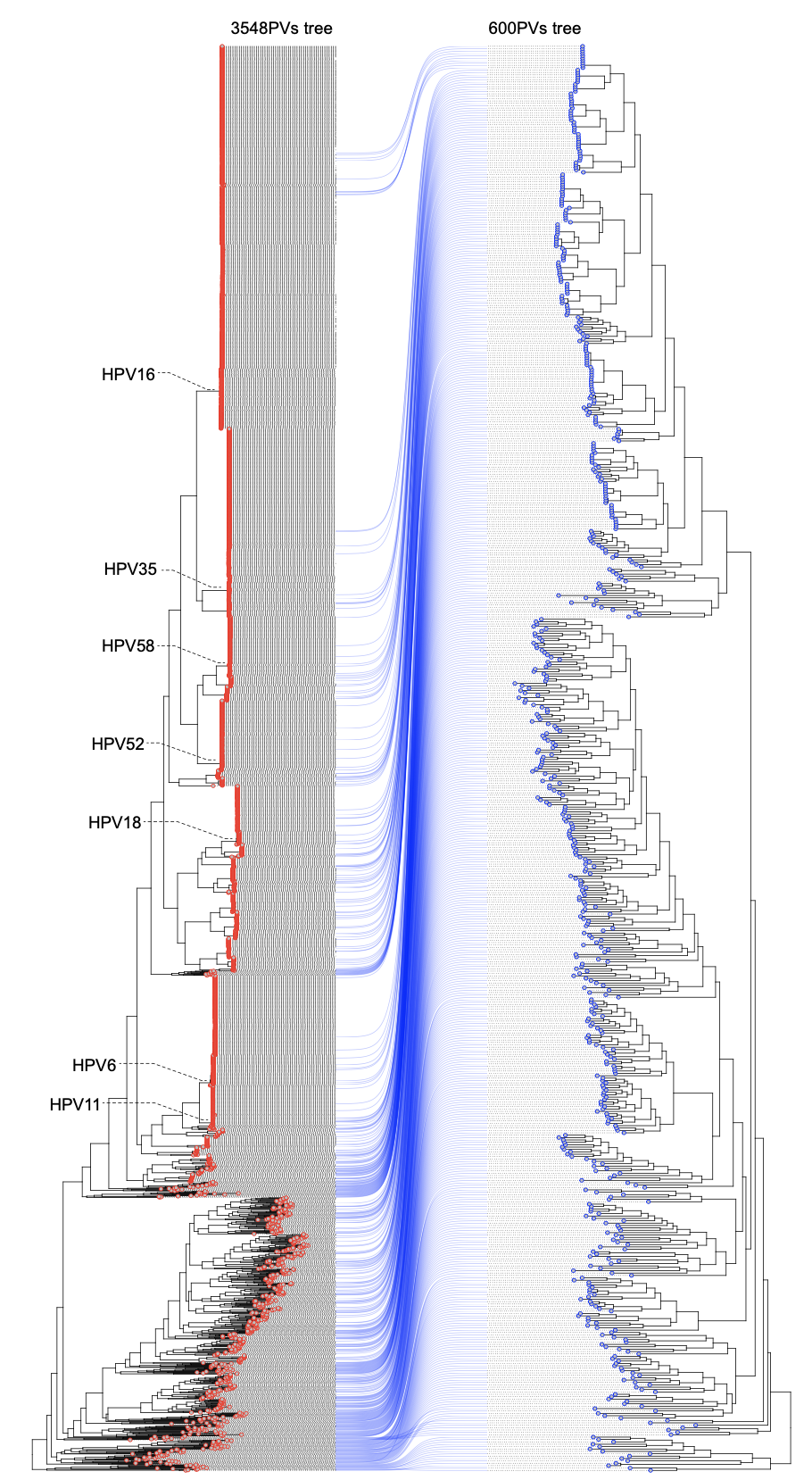

**Supplementary Figure 7. Phylogenetic correspondence between the 3548 PV tree and the downsampled 600 PV tree.**

Some representative HPV lineages in the 3548 PV tree is indicated.

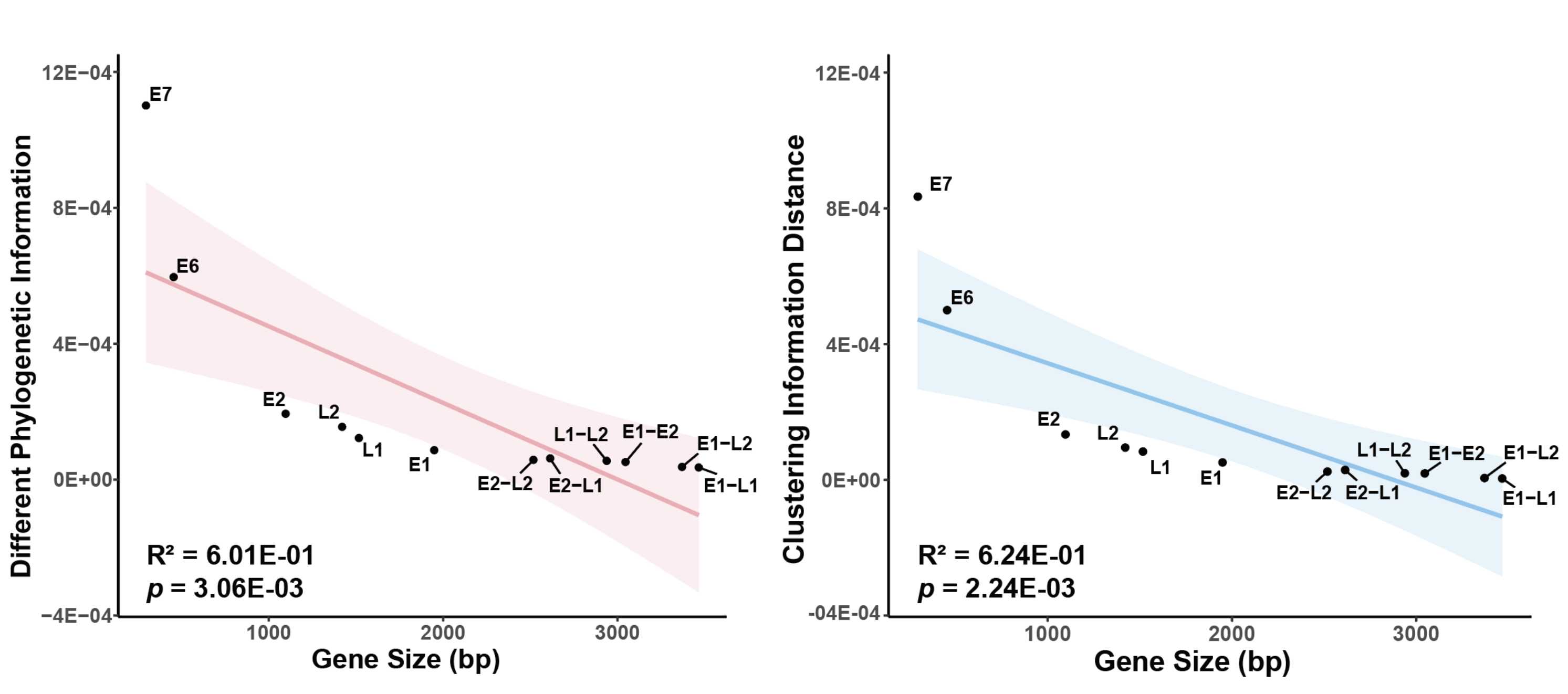

**Supplementary Figure 8. The correlation between phylogenetic distance and gene sizes based on 600 representative PV genomes.**

Different phylogenetic information (DPI; left panel) and clustering information distance (CID; right panel) were calculated to quantify their phylogenetic distance.

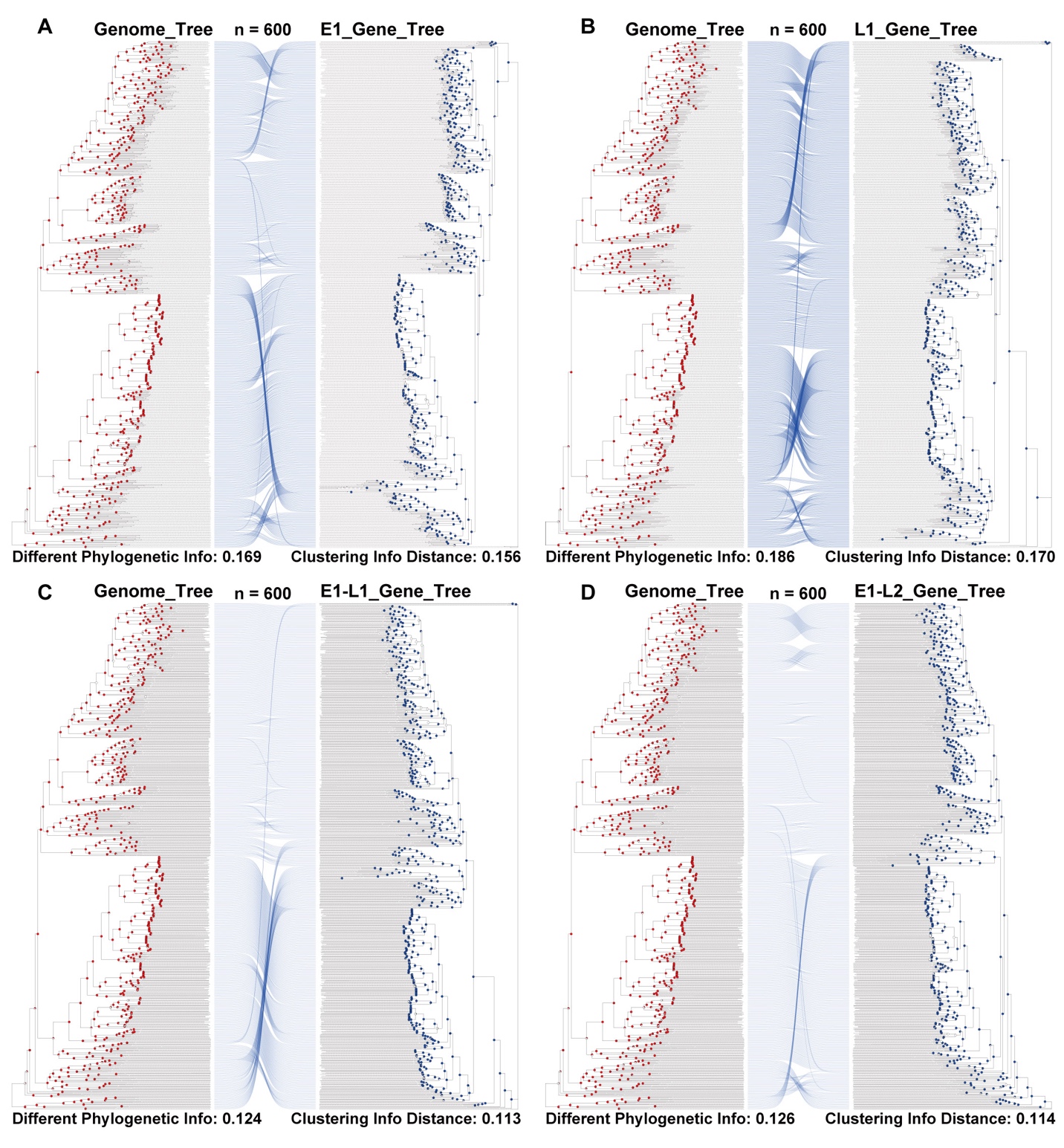

**Supplementary Figure 9. The genome tree and gene tree topology comparison based on 600 representative PV genomes.**

Gene trees derived from (A) E1, (B) L1, (C) E1-L1, (D) E1-L2 on the right were compared against the genome tree on the left. Different phylogenetic information (DPI) and clustering information distance (CID) were calculated to quantify their phylogenetic distance.

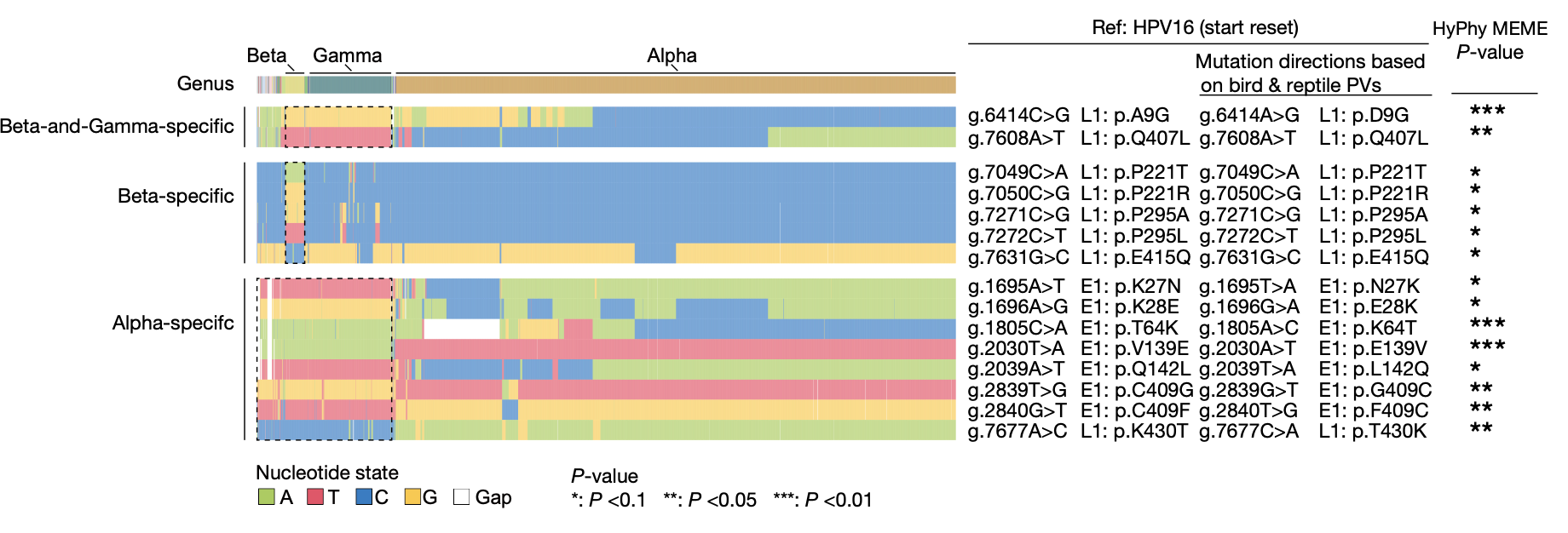

**Supplementary Figure 10. The nucleotide state composition at our identified lineage-defining PV variant sites.**

The genomic coordinates are based on the start-reset HPV16 reference genome. HyPhy MEME was used to score the significance of diversifying selection.

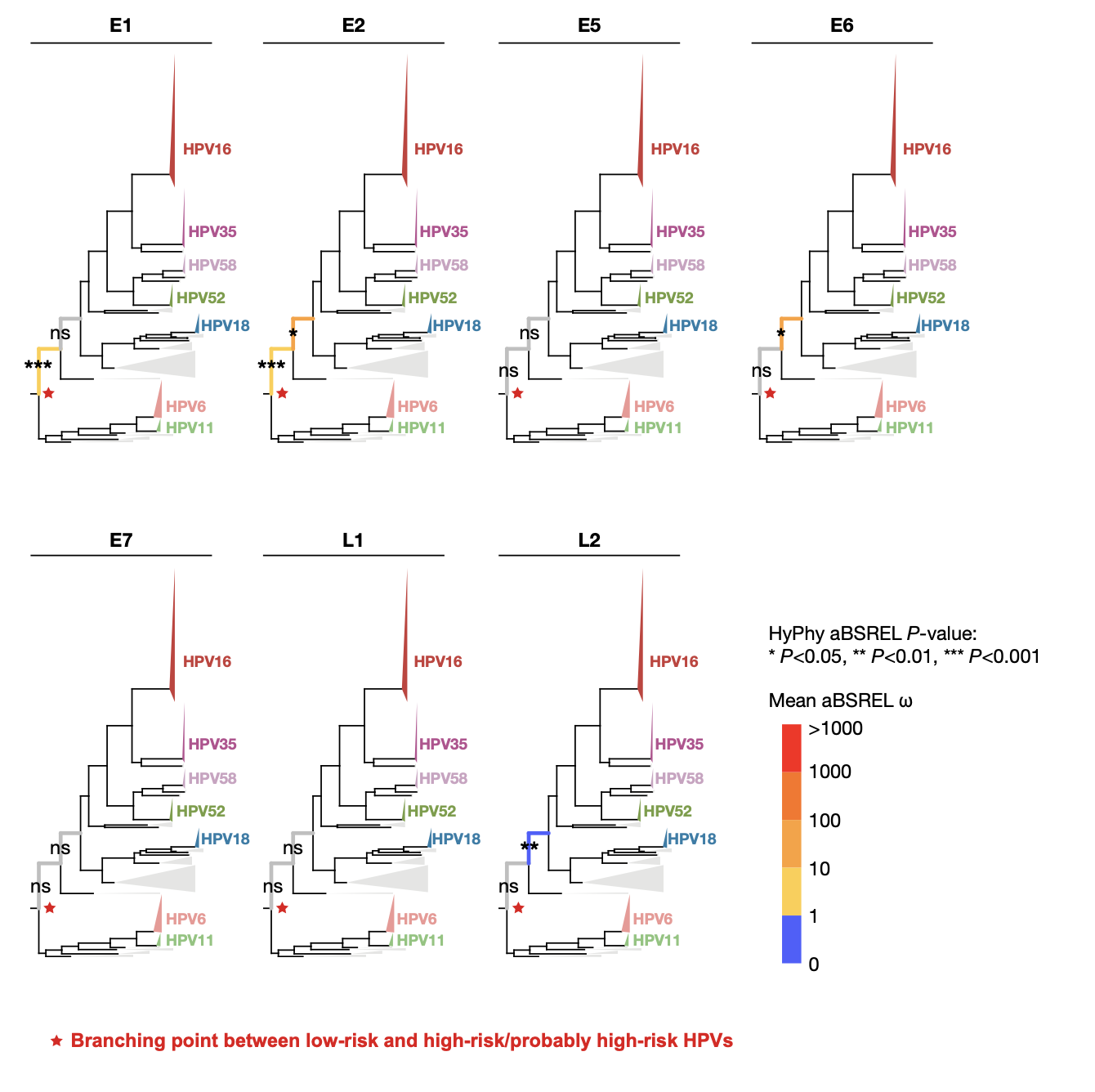

**Supplementary Figure 11. Branch-specific diversifying selection along the evolutionary divergence between low-risk and high-risk HPVs.**

The HyPhy aBSREL test was used to evaluate the significance of branch-specific diversifying selection immediately before and after the phylogenetic divergence point between low-risk and high-risk/probably high-risk HPVs.

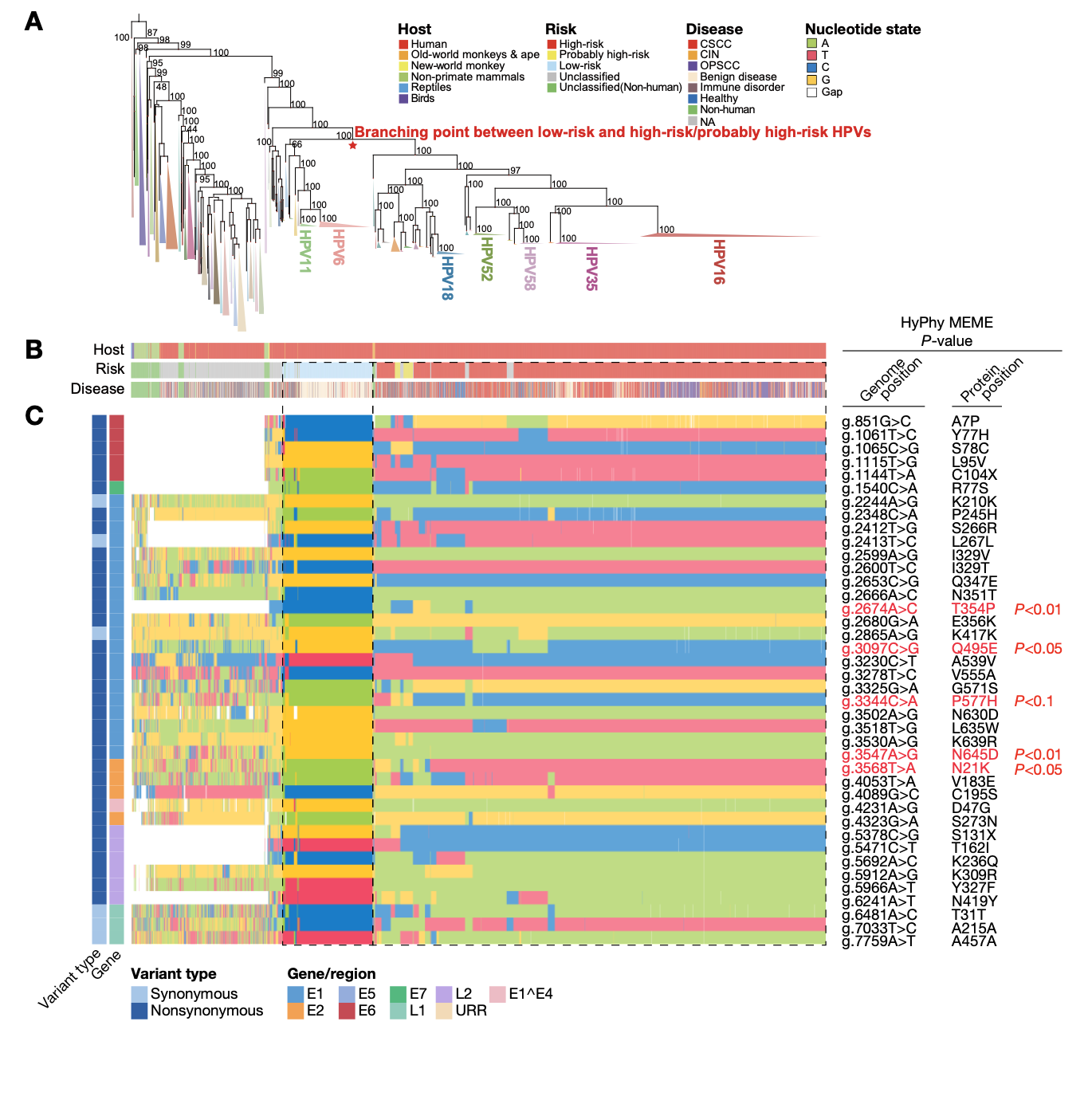

**Supplementary Figure 12. Overview of the 40 risk-defining variant sites regarding the evolutionary divergence between low-risk and high-risk HPVs.**

(A) The PV phylogenetic tree shown in Supplementary Figure 6. (B) Phylogenetic tree matched tracks corresponding to host species groups, risk classification, and disease types. (C) Phylogenetic tree matched nucleotide status pattern for the 40 risk-defining variant sites. The genomic coordinates of the variant sites are based on the start-reset HPV16 reference genome. The HyPhy MEME test was used to evaluate the significance of site-specific diversifying selection.

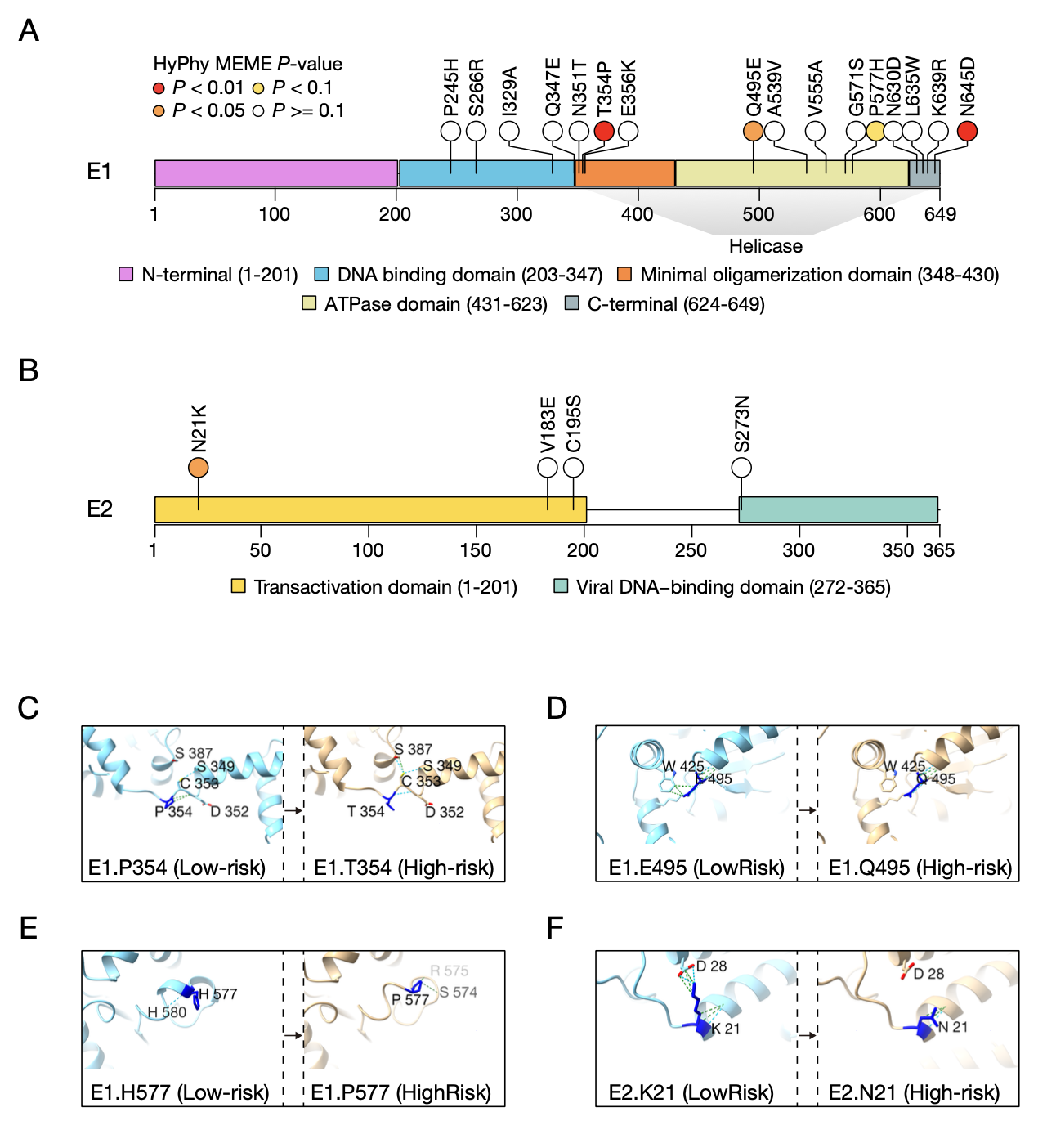

**Supplementary Figure 13. Distribution of our identified risk-defining variant sites in E1 and E2 proteins and their impacts on protein structure.**

(A, B) The distribution of risk-defining variant sites regarding the domain structure of E1 (A) and E2 (B) proteins. The site-specific diversifying selection was evaluated by HyPhy MEME test. (C-F) The functional impacts of the four selection-driven nonsynonymous risk-defining variant sites on the intramolecular interactions of their residing protein (i.e., E1 or E2).

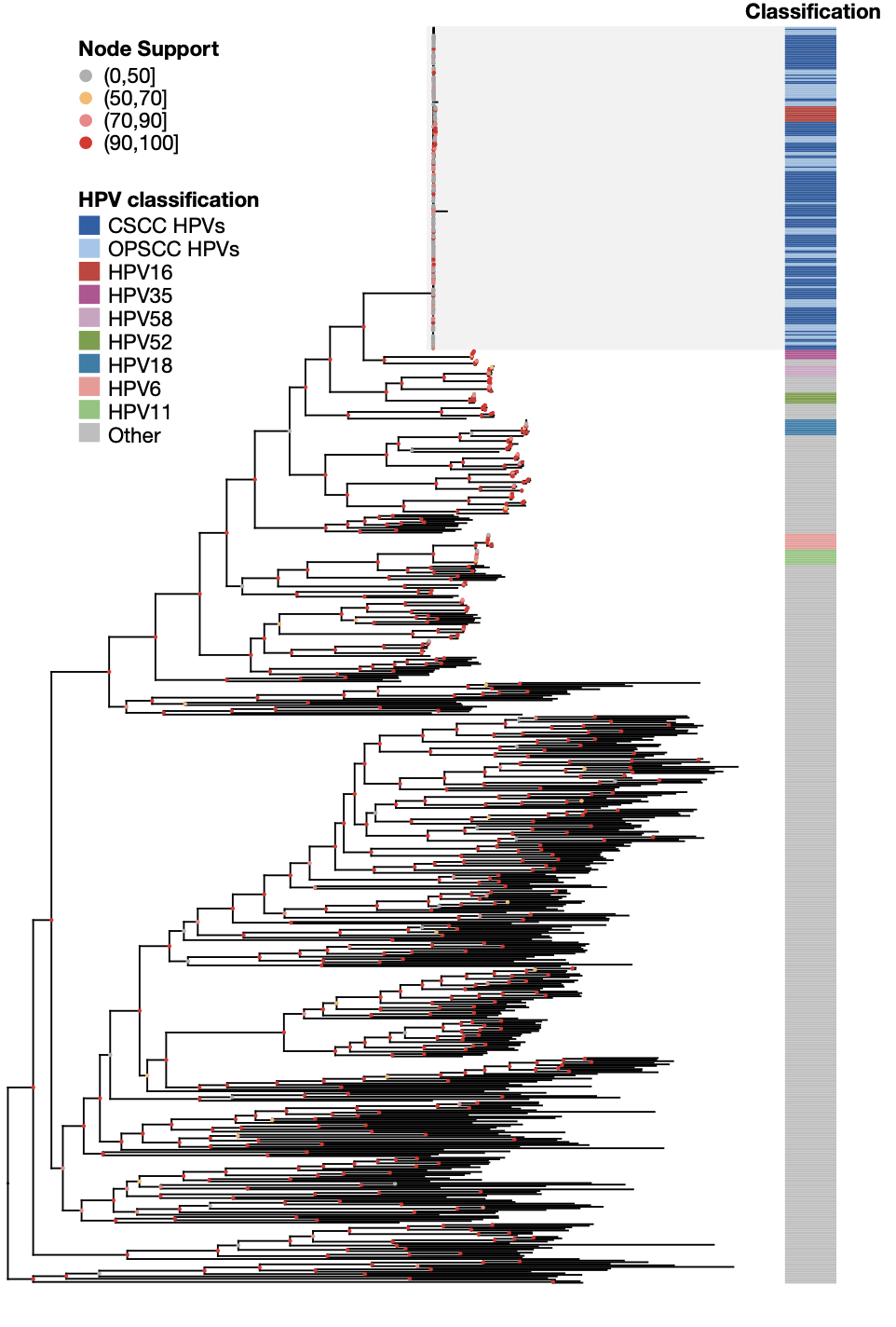

**Supplementary Figure 14. Phylogenetic positioning of our sequenced HPV genomes from CSCC and OPSCC patient cohorts relative to the 600-PV set.**

The labeled node support values were calculated by approximate likelihood-ratio test (aLRT).

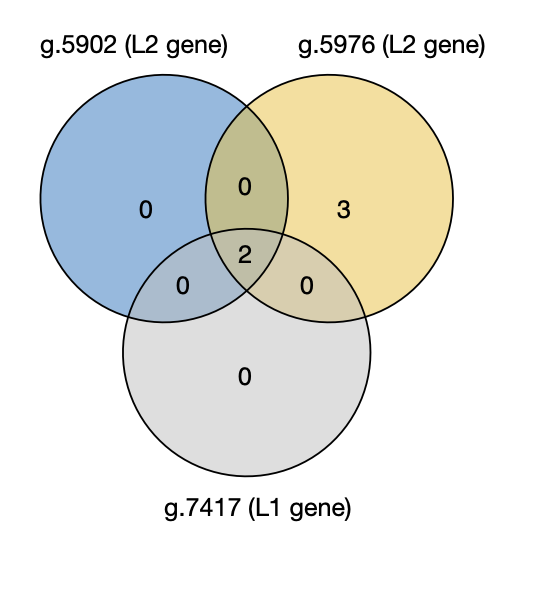

**Supplementary Figure 15. The number of overlapping samples for the three CSCC prognostic HPV16 variants.**

The genomic coordinates shown are based on HPV16 reference genome with start reset.

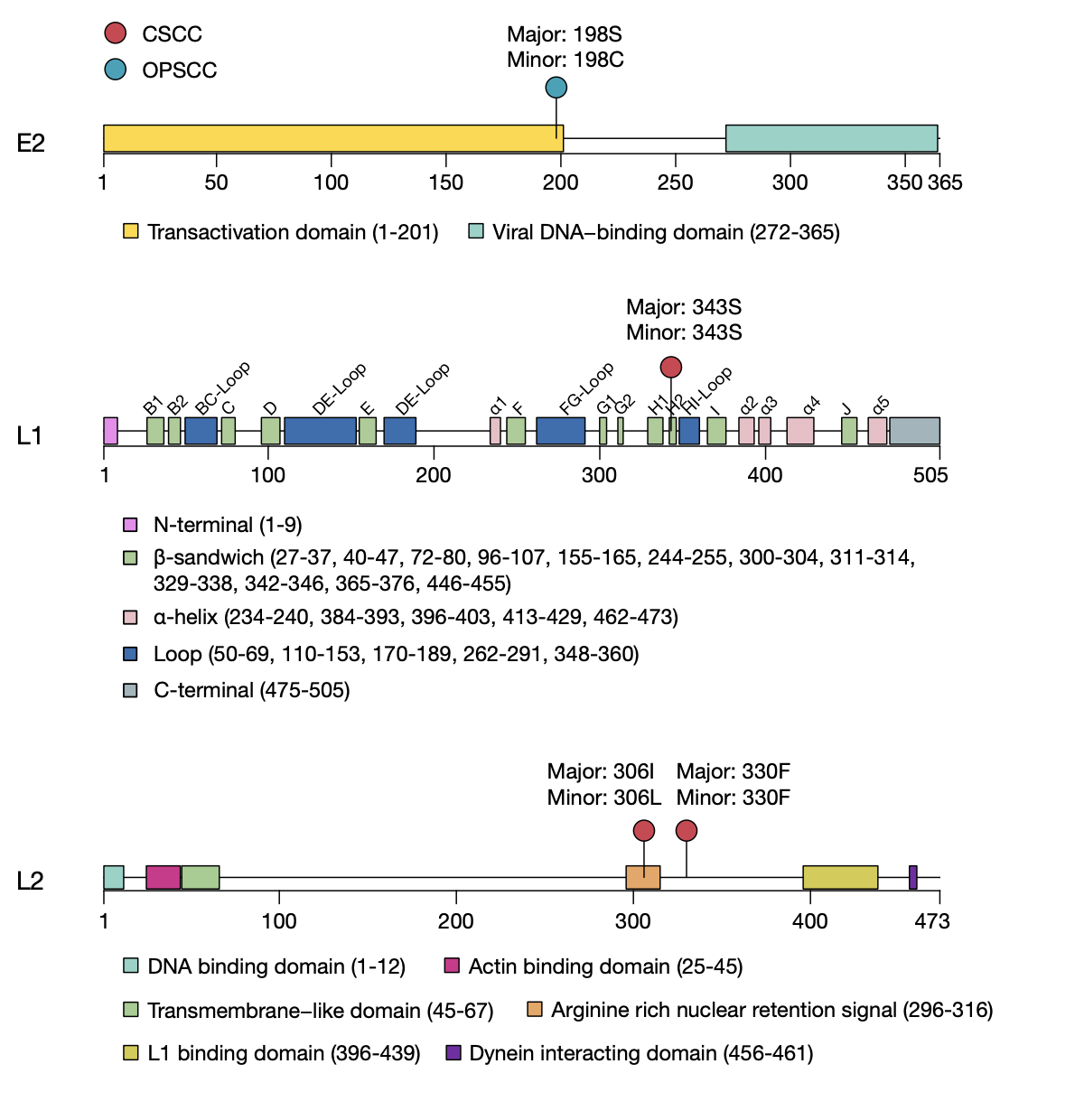

**Supplementary Figure 16. Distribution of our identified prognostic variant sites in the CSCC and OPSCC cohorts in their corresponding HPV16 protein space.**

The domain structures of the E2, L1, and L2 proteins were annotated based on literatures. The plotted minor allele is the risk-form regarding prognosis.

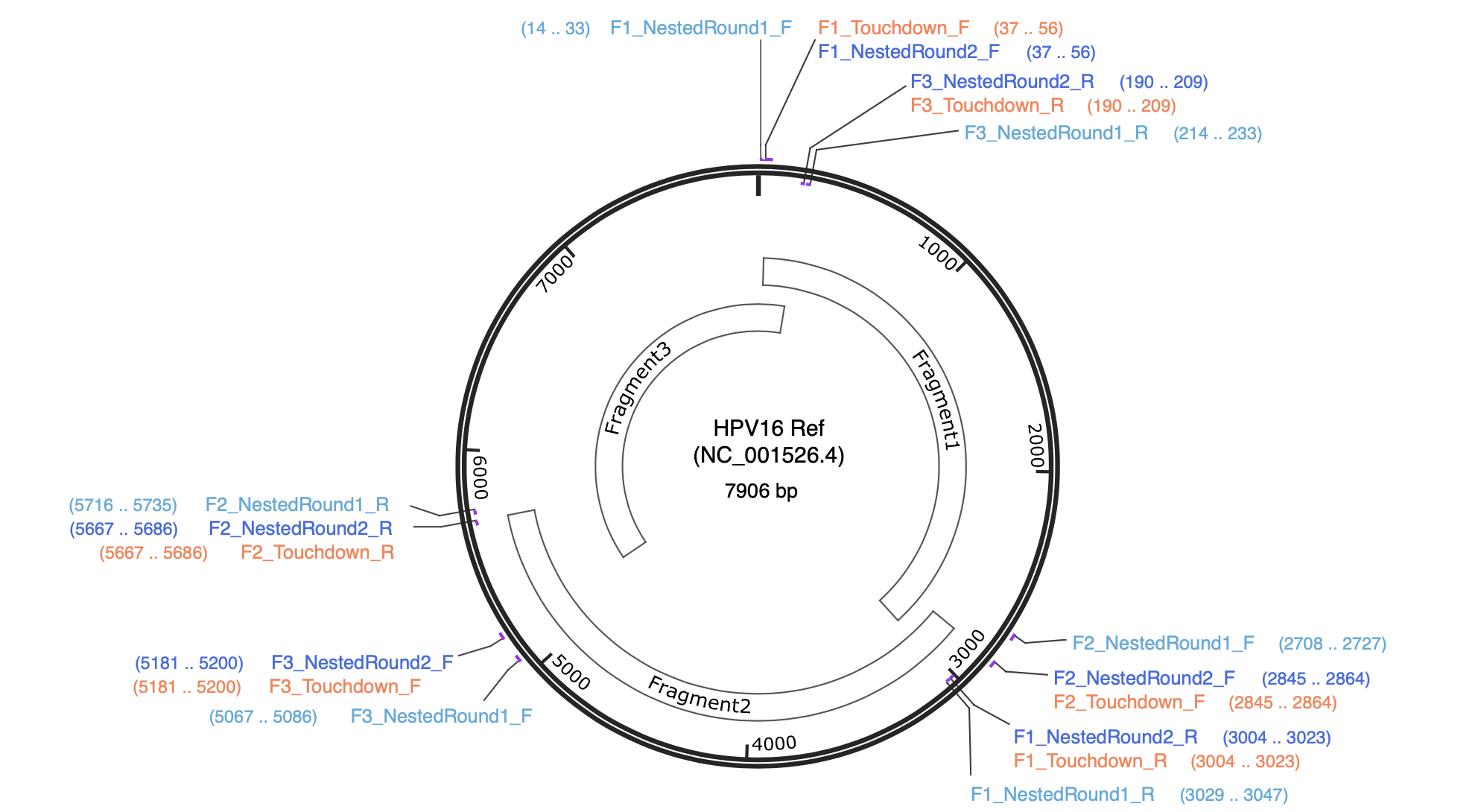

**Supplementary Figure 17. Primer design for HPV16 genome amplification.**

The primers used for touchdown PCR (TD-PCR) are labeled with “Touchdown”. The primers used for nested PCR were labeled with “Nested”. The forward and reverse primers are indicated by “F” and “R”.
